# Immunosuppressive traits of the hybrid epithelial/mesenchymal phenotype

**DOI:** 10.1101/2021.06.21.449285

**Authors:** Sarthak Sahoo, Sonali Priyadarshini Nayak, Kishore Hari, Prithu Purkait, Susmita Mandal, Akash Kishore, Herbert Levine, Mohit Kumar Jolly

**Affiliations:** Undergraduate program, Indian Institute of Science, Bangalore, Karnataka 560012, India; Centre for BioSystems Science and Engineering, Indian Institute of Science, Bangalore, Karnataka 560012, India; College for Integrated Studies, University of Hyderabad, Hyderabad, Telangana 500046, India; Department of Computer Science & Engineering, SSN College of Engineering, Chennai, India; Center for Theoretical Biological Physics, Northeastern University, Boston, MA; Departments of Physics and Bioengineering, Northeastern University, Boston, USA

**Keywords:** Hybrid epithelial/mesenchymal, PD-L1, immune evasion, multistability

## Abstract

Recent preclinical and clinical data suggests enhanced metastatic fitness of hybrid epithelial/ mesenchymal (E/M) phenotypes, but mechanistic details regarding their survival strategies during metastasis remain unclear. Here, we investigate immune-evasive strategies of hybrid E/M states. We construct and simulate the dynamics of a minimalistic regulatory network encompassing the known associations among regulators of EMT (epithelial-mesenchymal transition) and PD-L1, an established immune-suppressor. Our model simulations, integrated with single-cell and bulk RNA-seq data analysis, elucidate that hybrid E/M cells can have high levels of PD-L1, similar to those seen in cells with a full EMT phenotype, thus obviating the need for cancer cells to undergo a full EMT to be immune-evasive. Specifically, in breast cancer, we show the co-existence of hybrid E/M phenotypes, enhanced resistance to anti-estrogen therapy and increased PD-L1 levels. Our results underscore how the emergent dynamics of interconnected regulatory networks can coordinate different axes of cellular fitness during metastasis.

## Introduction

The progression of cancer relies on a complex interplay of various cell autonomous and non-cell autonomous phenomena. The latter includes the well-established fact that cancer cells can proactively create a microenvironment that aids their own survival. One of the employed strategies is to suppress various arms of immune system that can lead to cancer cell elimination (1). For instance, some tumor cells can inhibit the functions of effector T (T_eff_) cells and/or induce a population of tolerogenic cells that ultimately result in the immune escape of the tumor. They can also facilitate accumulation of immune suppressive cells such as regulatory T (T_reg_) cells, myeloid derived suppressor cells (MDSCs) and M2 macrophages/ tumor-associated macrophages (TAMs), leading to enhanced tumor growth (1). Understanding these strategies of tumor-driven reprogramming of the microenvironment would be a major step towards more effective guiding of various therapeutic interventions.

In addition to reprogramming the immune cells in the stroma, tumors employ cell autonomous mechanisms that help them directly evade cytotoxic CD8 T cells. A key mechanism via which tumor cells achieve this evasion is via the expression of programmed death-ligand 1 transmembrane protein (PD-L1) on their cell membranes (2). The binding of PD-L1 to PD-1 receptors on activated T cells drives the exhaustion of these T cells, reducing their cytotoxic abilities (3). In cancer cells, a multitude of molecular players modulate PD-L1 levels at various regulatory stages (2). Of interest here is the finding that PD-L1 levels can be increased as cells go through an Epithelial-Mesenchymal Transition (EMT) and consequently gain the ability to migrate and invade (4–6). The process of EMT, however, is not typically a binary switch, as had been tacitly assumed in these earlier works. Instead, cells can stably maintain one or more hybrid epithelial/mesenchymal (E/M) phenotypes that can be much more metastatic than cells in a ‘full EMT’ or ‘extremely mesenchymal’ state (7). Besides, hybrid E/M phenotypes across cancers can be resilient to various therapies (7). However, the immune evasive properties of the hybrid E/M states are relatively poorly understood.

In this study, we identify a core regulatory network that helps us elucidate the immune evasive properties of different phenotypes along the epithelial-hybrid-mesenchymal spectrum. Our simulations indicate that hybrid E/M phenotypes are extremely likely to exhibit high PD-L1 levels, similar to those seen in mesenchymal cells, thus obviating the need to undergo a full EMT to develop immunosuppression. Moreover, the switch from an epithelial/low-PDL1 state to hybrid/ high-PDL1 or mesenchymal/high-PDL1 state is reversible, i.e., while EMT can induce PD-L1 levels, MET can reduce them. Specifically, for breast cancer we show how acquisition of reversible resistance to targeted therapy such as tamoxifen can co-occur with high PD-L1 levels, thus enabling cross-resistance and enhancing cancer cell fitness during metastasis. Our model predictions are validated by extensive analysis of transcriptomic datasets across multiple cancers at both bulk and single cell levels.

## Results

### Hybrid E/M and Mesenchymal cell states are more likely to exhibit high PD-L1 levels

Capturing the essence of biological processes via mechanism-based mathematical modelling can be a daunting task, given the vast complexity of biological systems. Identifying an appropriately sized gene regulatory network that incorporates the essential features of the underlying biological mechanism at hand in a minimalistic, yet informative manner is a key first step. To that extent, we started out with a core set of four well-reported biomolecules and their interactions that can capture the non-binary nature of the epithelial-mesenchymal plasticity (EMP) spectrum and the ability of different phenotypes to modulate PD-L1 levels: ZEB1, miR-200, CDH1 (E-cadherin) and SLUG (**Fig. 1A**). Mutual inhibition between ZEB1 and miR-200, together with ZEB1 self-activation can enable epithelial, mesenchymal and hybrid epithelial/mesenchymal (E/M) phenotypes (8). SLUG has been reported to specifically associate with hybrid E/M phenotype both in experimental and computational analysis (9,10).

**Fig 1.**
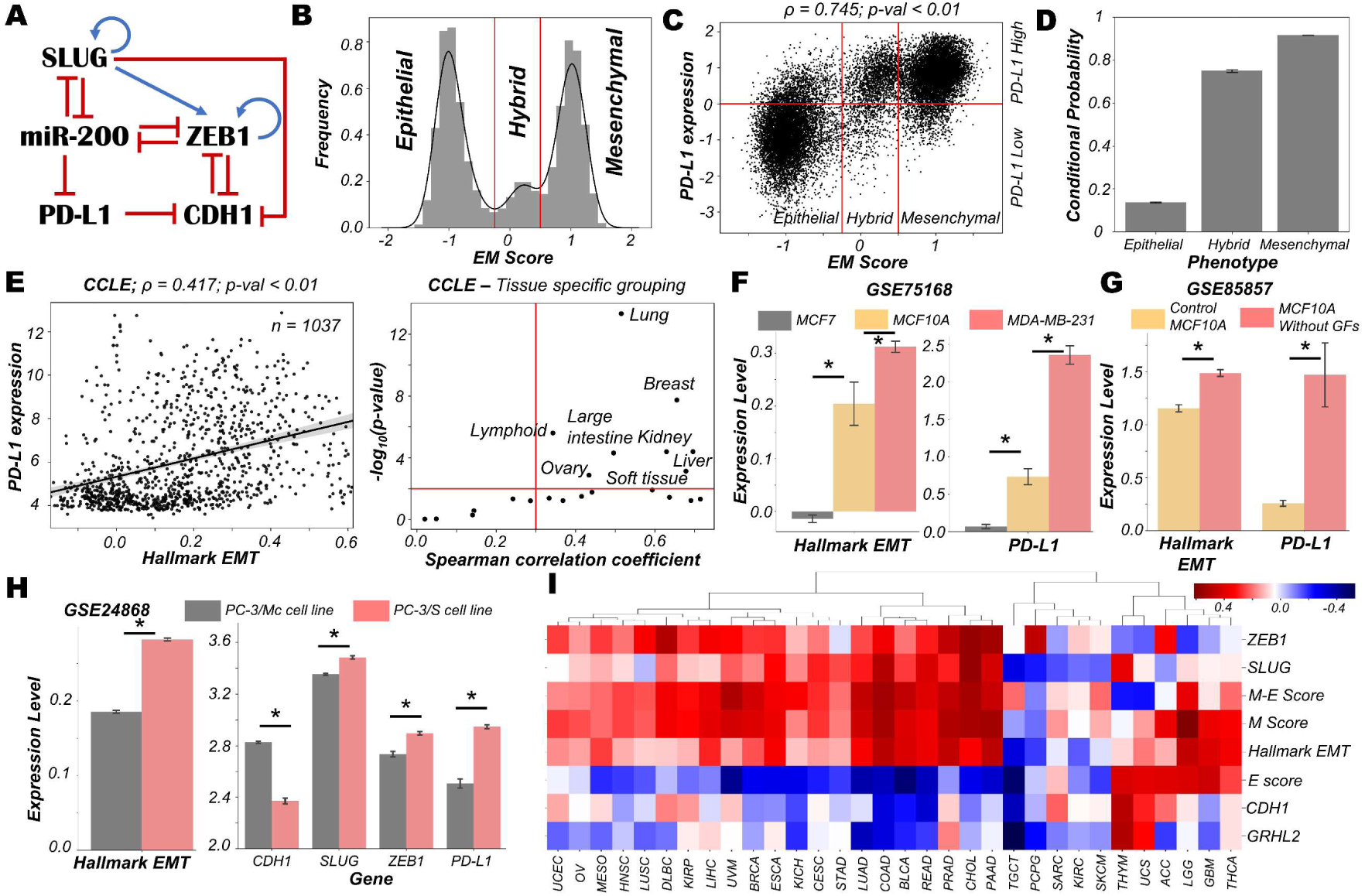
Dynamics of regulatory network coupling EMT with PD-L1. **A)** Regulatory network (GRN) capturing the interplay of EMT regulators coupled with PD-L1. Blue arrows stand for activation links, red hammers for inhibitory links. **B)** Density histogram of EM Score fitted with kernel density estimate showing a trimodal distribution. Red lines show the partition between phenotypes: Epithelial, Hybrid, and Mesenchymal. **C)** Scatter plot of PD-L1 expression and EM score. Horizontal red line shows the partition between PDL1 expression level being high vs. low. Vertical red lines show the partition between phenotypes: Epithelial, Hybrid, and Mesenchymal based on EM score. Spearman’s correlation coefficient (ρ) and corresponding p-value (p-val) have been reported. **D)** Bar plot representing the conditional probability of a phenotype being PD-L1 high given that it belongs to a given EMT phenotype. **E)** Scatter plot showing correlation between PD-L1 levels and the Hallmark EMT signature in cell lines from CCLE. Spearman’s correlation coefficient (ρ) and corresponding p-value (p-val) are reported (left panel). Splitting CCLE cell lines reveals tissues that show a strong significant correlation (ρ > 0.3 and p-val < 0.01) (right panel). **F)** Activity levels of Hallmark EMT and PD-L1 expression levels in 3 breast cancer cell lines (GSE75168). **G)** Activity/Expression levels of Hallmark EMT and PD-L1 levels in MCF10A breast cancer cells treated with or without growth factors (GSE85857). **H)** Activity/Expression levels of Hallmark EMT and PD-L1 levels in two prostate cancer sub-lines of PC3 with different EMT status (GSE24868). **I)** Heatmap showing the Spearman’s correlation coefficients between the different EM metrics and EMT associated genes and PD-L1 levels across 32 different cancer types in TCGA. * denotes a statistically significant difference (p-val < 0.05) between the represented groups assessed by a two-tailed Students t-test assuming unequal variances.

High levels of ZEB1 and SLUG with low levels of CDH1 and miR-200 are frequently attributed to a mesenchymal phenotype, while high levels of CDH1 and miR-200 with concurrent reduced levels of ZEB1 and SLUG usually associates with an epithelial phenotype (11). Interactions between ZEB1, miR-200, CDH1, and SLUG have been extensively studied (10–13). Furthermore, miR-200 has been known to directly inhibit PD-L1 by binding to 3’ untranslated region of its mRNA (4). PD-L1 can, in turn, repress the levels of CDH1 via indirect mechanisms (14) (**Fig 1A**).

We used RACIPE (15) to generate *in-silico* steady state gene expression values enabled by this gene regulatory network (**Methods**). RACIPE simulates a given gene regulatory network as a set of coupled ordinary differential equations (ODEs), with parameters sampled from biologically relevant ranges. The ensemble of resultant steady states is indicative of the possibility space allowed by the network topology. To quantify the cellular phenotype of given steady state solution, we defined an EM score from z-normalised expression values of ZEB1, SLUG, miR-200, and CDH1. The higher the EM score, the more mesenchymal is the corresponding phenotype. A histogram of these scores showed a clear trimodal distribution, which can be construed as consisting of epithelial, hybrid E/M, and mesenchymal phenotypes; these assignments can be confirmed by PCA plots (**Fig 1B, S1A-B, Table S1**). Subsequently, we also observed a bimodal distribution of PD-L1 levels (**Fig S1C**) where high levels of PD-L1 can be viewed as an immune-evasive state while low PD-L1 denotes an immune-sensitive tumor cell state (16).

Next, we investigated the association between the EM scores and PD-L1 levels and observed a strong positive correlation between them (ρ = 0.745; p-value < 0.01) (**Fig 1C**). Conditional probability analysis shows that only a small percentage (∼15%) of epithelial cells were PD-L1+. In contrast, a much larger percentage of hybrid E/M (∼70%) and mesenchymal (∼90%) cells were PD-L1+ (**Fig 1D, S1D**). Consistently, PD-L1 was found to negatively correlate with CDH1 but positively with ZEB1 and SLUG (**Fig S1E**). Further, ZEB1, SLUG, and PD-L1 all had intermediate levels in hybrid E/M states compared to extreme states – epithelial and mesenchymal (**Fig S1F**). Together, these results suggest that cells undergoing either a partial or full EMT can upregulate their levels of PD-L1 and consequently can exhibit immune evasion.

To validate these model predictions, we analysed pan-cancer gene expression datasets such as CCLE (Cancer Cell Line Encyclopaedia), where we observed the ssGSEA scores of EMT to be positively correlated with PD-L1 levels (**Fig 1E; left**). A tissue-specific analysis revealed a majority (16 out of 22) of cancers exhibited a strong correlation (ρ > 0.3) between EMT and PD-L1 expression (**Fig 1E; right, Table S2**). Next, we investigated more specific scenarios. For instance, three breast cancer cell lines along the EMP spectrum – MCF7 (epithelial), MCF10A (hybrid E/M) and MDA-MB-231 (mesenchymal) (17,18) – showed consistent trends with PD-L1 levels with MCF7 < MCF10A < MDA-MB-231 (**Fig 1F**). This pattern was recapitulated in an analysis of TCGA luminal breast cancer samples (**Fig S1G**). Furthermore, MCF10A cells, when driven to a more mesenchymal phenotype upon growth factor depletion (19), showed a concomitant increase in the levels of PD-L1 (**Fig 1G**). Similarly, comparing two sub-lines of prostate cancer cells PC-3 (20), the more mesenchymal one (PC-3/S) showed higher levels of PD-L1, ZEB1, and SLUG relative to the hybrid E/M PC-3/Mc cells (**Fig 1H**). A positive correlation between EMT signature and PD-L1 levels was also seen in A549 lung adenocarcinoma cells induced to undergo EMT (**Fig S1H**), suggesting a pan-cancer association of EMT with PD-L1 levels.

Finally, we analysed TCGA patient cohort datasets for all the above-mentioned features. We calculated the Spearman’s correlation coefficient between PD-L1 expression levels with those of CDH1, GRHL2, SLUG, and ZEB1 as well as with a Hallmark EMT signature, and epithelial and mesenchymal specific signatures (**Fig 1I, S2A**). A majority of cancers (21 out of 32) showed a strong positive correlation of PD-L1 with the mesenchymal related metrics (ZEB1, SLUG, Hallmark EMT, mesenchymal score, and M-E score) while showing an intermediate to strong negative correlation with the epithelial ones (CDH1, GRHL2, and epithelial score), thereby endorsing our model predictions. Intriguingly, 6 cancer types (THYM, UCS, ACC, LGG, GBM, and THCA) showed a positive correlation of PD-L1 with both epithelial and mesenchymal signatures, highlighting a possible association of highest PD-L1 levels with the hybrid E/M phenotype. It should be noted that our model does not completely preclude the association of an epithelial state with high PD-L1 levels, although the likelihood of such association is relatively low (**Fig 1C**). This infrequent association may underlie context-specific behavior of epithelial tumours (such as Thymic epithelial tumours) that also can show high PD-L1 positivity (**Fig 1I**) (21).

Overall, *in silico* predictions, supported by analysis of *in vitro* and patient data, suggests that a change in the EMP status of the cell is positively associated with PD-L1 levels across various cancer types. These results clearly indicate the likelihood of the hybrid E/M phenotype being (almost) as immune evasive as the mesenchymal phenotype.

### Traversal of cells on the EMP spectrum can alter the PD-L1 status of the cells

After establishing a pan-cancer correlation between more mesenchymal status and higher PD-L1 levels, we examined a causal connection between them. We simulated the set of coupled ODEs for a representative model whose parameter set gave rise to tristability. We started with diverse initial conditions and observed convergence to three distinct EM states (**Fig. 2A**). Corresponding PD-L1 levels followed the previously observed patterns with epithelial state showing the least PD-L1 levels, while both the hybrid E/M and mesenchymal showing nearly equal levels of PD-L1 higher than those of the epithelial state (**Fig 2A**). Stochastic simulations for this parameter set created a landscape indicative of the steady states under the influence of biological noise. This landscape revealed the co-existence of three states (shown by valleys) – (Epithelial, PD-L1 low), (Hybrid E/M, PD-L1 high) and (Mesenchymal, PD-L1 high) (**Fig. 2B**), depicting that as cells change their EM status, their corresponding PD-L1 levels are also altered. In another parameter set to study the stochastic dynamics of this network (22), we observed spontaneous switches between epithelial and mesenchymal states with a concurrent change in the levels of PD-L1 (**Fig. 2C**). Together, this analysis points towards the possibility of a switch like behaviour in acquisition of an immune evasive phenotype as the cells undergo EMT.

**Fig 2.**
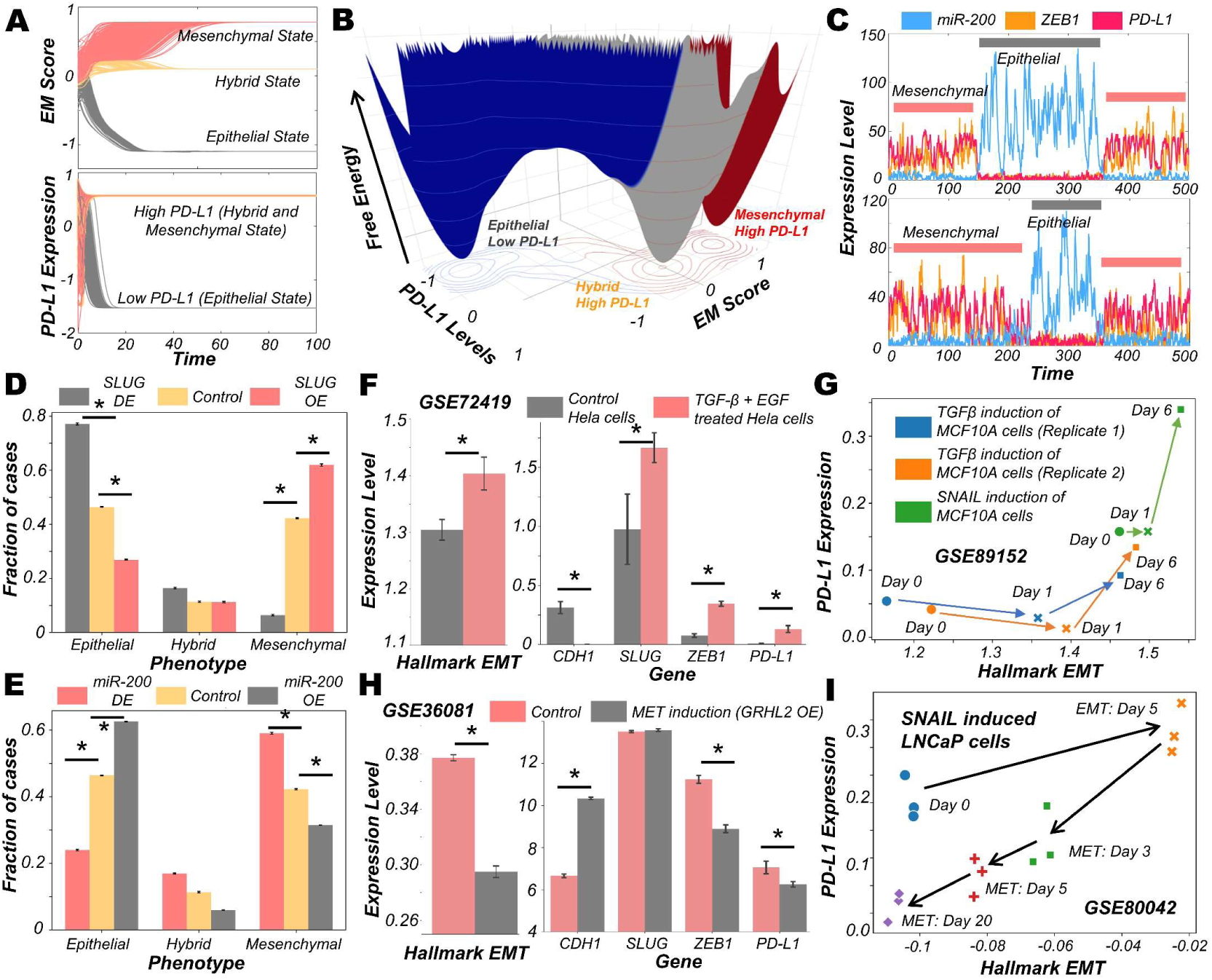
Evidence for causal links between EMT associated genes and PD-L1 levels. **A)** Dynamics of EM score and PD-L1 showing presence of epithelial, hybrid, and mesenchymal phenotypes and their corresponding PD-L1 levels, when simulated from multiple initial conditions. **B)** Probability landscape on the PD-L1 and EM score plane, with the valleys representing the stable states possible in the system. Three distinct states – Epithelial/ PD-L1 low, Hybrid E-M/ PD-L1 high, and Mesenchymal/ PD-L1 high – are observed. **C)** Stochastic simulations of gene regulatory network via sRACIPE showing spontaneous switching between different states. **D)** Simulation results showing the fraction of cases of epithelial, hybrid, and mesenchymal phenotypes under control (yellow), SLUG DE (grey) and SLUG OE (orange) conditions. **E)** Same as D but for miR-200 OE (grey) and miR-200 OE (orange) conditions. **F)** Activity/Expression levels of Hallmark EMT and PD-L1 levels in Hela cells induced to undergo EMT (GSE72419). **G)** Two-dimensional Hallmark EMT and PD-L1 plot showing trajectory of MCF10A cells induced with TGFβ or SNAIL (GSE89152). **H)** Activity/Expression levels of Hallmark EMT and PD-L1 levels in HMLE cells where MET has been induced via overexpression of GRHL2 (GSE36081). **I)** Two-dimensional Hallmark EMT and PD-L1 plot showing trajectory of LNCaP prostate cancer cells that have been induced with SNAIL to undergo EMT followed by removal of signal to induce MET (GSE80042). * denotes p-val < 0.05 as assessed by a two-tailed Students t-test assuming unequal variances.

To characterize the impact of perturbations on our core regulatory network we simulated the scenarios of EMT induction and MET induction. EMT was induced by down expression (DE) of miR-200 or over expression (OE) of SLUG; conversely, MET was induced by OE of miR-200 or DE of SLUG (13). SLUG-OE or miR-200 DE increased the proportion of mesenchymal cell states with a concurrent decrease in epithelial cases (**Fig 2D-E, Fig S3A-B, D-E**). This change resulted in a significant increase in PD-L1 levels (**Fig S3C, F**). Opposite trends were observed in the cases of MET induction via miR-200 or SLUG-DE, with resultant changes in PD-L1 levels (**Fig 2D-E**).

Next, we probed whether our model prediction about a concurrent switch in EM status and PD-L1 levels is supported by experiment, through analysing corresponding transcriptomic data. HeLa cells treated with TGFβ and EGF were thereby induced to undergo EMT, evident by increases in SLUG and ZEB1 levels, as well in the activity (as measured via ssGSEA; see Methods) of the Hallmark EMT gene set (**Fig 2E**). In treated cells, CDH1 levels were significantly decreased while PD-L1 levels were increased (**Fig 2E**). This phenomenon of EMT-driven increase in PD-L1 was also seen in non-cancerous cells where TGFβ treatment of primary airway epithelial cells led to upregulation of EMT and PD-L1 (**Fig S3G**), indicating that this association between EMT and PD-L1 levels need not be restricted to cancer cells. Furthermore, we compared the profiles of triple negative breast cancer cells DKAT when grown in a medium supporting epithelial growth (MEGM) vs when grown in a medium containing stromal factors (SCGM). These have been shown to differ in their EM status: while culturing in SCGM facilitated a mesenchymal phenotype, that in MEGM drove an epithelial one (GSE33146). Consistently, SLUG, ZEB1 and PD-L1 levels were significantly higher in cells grown in SCGM rather than in MEGM (**Fig S3G**). Furthermore, in a time course experiment where EMT was induced in A549 lung cancer cells by treatment with TGFβ over 96 hours, the failure of sustained expression of ZEB1 was correlated with a visibly lower level of PD-L1 levels, hinting towards a likely causal role of ZEB1 in enhanced PD-L1 expression levels. (**Fig S3I**). Next, we analysed a set of experiments in which MCF10A cells were induced to undergo EMT either via TGFβ application or by induced overexpression of SNAIL. This time course experiment resulted in an increase of activity of Hallmark EMT genes and PD-L1, irrespective of cells’ initial EM status (**Fig 2F**).

Finally, we asked whether induction of MET can decrease the levels of PD-L1 in cancer cells. HMLE cells, upon overexpression of MET-inducing factor GRHL2, displayed a more epithelial state (increased CDH1, decreased ZEB1 and decreased hallmark EMT signature) with a substantial drop seen in PD-L1 levels (**Fig 2H**), indicating that EMT-driven changes in PD-L1 levels are reversible. Similar observations were made for PD-L1 levels in LNCaP prostate cancer cells which were first induced to undergo EMT and subsequently were induced to undergo MET. A two-dimensional plot of EM score and PD-L1 levels revealed an increase in PD-L1 as EMT was induced, and a subsequent decrease when MET was induced **(Fig 2I)**. Intriguingly, the cell population did not retrace its original path during MET induction, indicative of hysteresis in the system (23). The overall levels of PD-L1 were lower at the end of 20 days of MET than for the uninduced cells themselves, suggesting that MET induction can reset the baseline PD-L1 levels upon a cycle of EMT and MET. Collectively, these results underscore that induction of EMT or MET in cancer cells (and possibly other cells as well) can regulate their immune evasion status through altered levels of PD-L1.

### Various signalling pathways can either independently or in concert modulate the immune evasive properties of cancer cells on the EMP spectrum

The above-mentioned interconnections among the EMT regulators and PD-L1 levels seldom work in isolation. Multiple signalling pathways can independently or in concert affect the EM status of cells and/or their PD-L1 expression. To investigate such effects, we calculated the degree of correlation of 15 well-defined signalling pathways with EMT and with PD-L1 levels across different cancers in the TCGA cohort (**Fig S4A**). A scatter plot of corresponding correlation coefficients revealed pan-cancer consistency in signalling pathways associated with EMT and with PD-L1 levels (*ρ* = 0.37; p-value < 0.01) (**Fig S4B**). Next, we ranked which pathways correlate most strongly with EMT signature or PD-L1 levels (**Fig 3A**). TGFβ, IL2-STAT5, TNFα-NFκB, IL6-STAT3 and NOTCH signalling were found to correlate strongly with EMT, consistent with their expected roles (24). Similarly, PD-L1 levels are most correlated with IL6-STAT3, IFNγ, IL2-STAT5, IFNα and TNFα-NFκB, all of which have been previously implicated (25). Plotting these pathways through their normalised ranks allows identifying the pan-cancer independent regulators of PD-L1 levels and EMT; for instance, TGFβ and NOTCH can be considered as more EMT-specific, while IFNγ and IFNα are PD-L1 specific modulators. IL6-JAK-STAT3, IL2-STAT5 and TNFα-NFκB pathways correlated both with PD-L1 and EMT (**Fig 3B**). IL1β is known to act partially by the NFκB pathway (26). Thus, it is not surprising to see that treatment of cancer cells with IL1β caused a concerted increase in EMT as well as in PD-L1 levels; the consistency in these trends was also visible upon withdrawal of the signal (**Fig 3C**).

**Fig 3.**
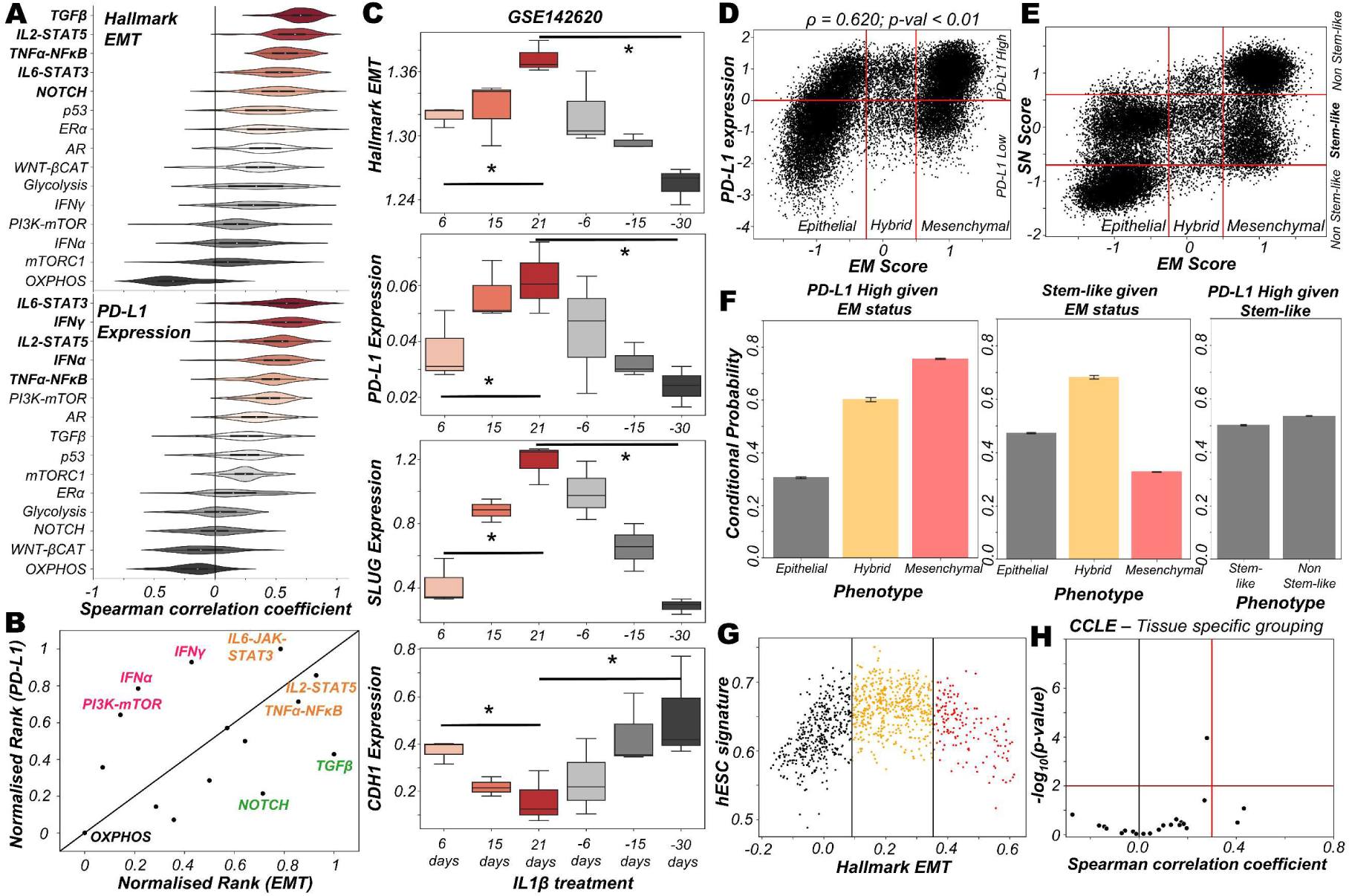
Signalling pathways and biological processes that can affect PD-L1 and/or EMT. **A)** Violin plots of Spearman’s correlation values of different signalling pathways with Hallmark EMT programme (top) or with PD-L1 levels (bottom) ordered by corresponding median values across 27 cancer types in TCGA. **(B)** Scattered plot between normalised ranks of signalling pathways with the EMT programme and with PD-L1 expression levels. Signalling pathways hypothesised to be specific to EMT programme are labelled in green, those specific for PD-L1 highlighted in pink and those with both in orange. **C)** Activity/expression levels of Hallmark EMT, PD-L1, SLUG, and CDH1 levels in lung cancer cells treated with IL-1β and subsequent removal of signal (GSE142620). **D)** Scatter plot of PD-L1 expression and EM score. Horizontal red line shows the partition between PD-L1 expression level being high vs low for the circuit in Fig S4C. Vertical red lines show the partition between phenotypes: Epithelial, Hybrid, and Mesenchymal based on EM score. Spearman’s correlation coefficient (ρ) and corresponding p-value (p-val) are reported. **E)** Scatter plot of SN score and EM score showing the presence of clusters having predominantly stem-like hybrid and the presence of both stem like and non-stem like epithelial and mesenchymal cells scattered in the plane. Horizontal red lines show the partition between stem-like and non-stem-like based on SN score and EM phenotypes. **F)** Bar plot representing conditional probability of PD-L1 being high given EM status, stem-like phenotype given EM status, and PD-L1 high given stemness status respectively. **G)** Scatter plot showing the non-monotonic association between the hESC signature and the Hallmark EMT signature in CCLE dataset. The boundaries are determined by trisection of the entire range of Hallmark EMT signature values. **H)** No tissue in CCLE shows a strong significant Spearman’s correlation (ρ > 0.3 and p-val < 0.01) between hESC signature and PD-L1 levels. * denotes a statistically significant difference (p-val < 0.05) between the represented groups assessed by a two-tailed Students t-test assuming unequal variances.

The interplay between stemness and EMT has been extensively investigated (27,28). Thus, we asked whether EMT, stemness and PD-L1 levels all vary together. To investigate this crosstalk, we simulated an extended regulatory network including stemness regulators (OCT4, miR-145, LIN28, let-7) via RACIPE (**Fig S4C)**. A stemness window was defined based on the distribution of stemness (SN) score (**Fig S4D**). This network showed conserved trends between EM score and PD-L1 expression level and found that most hybrid E/M solutions lay within the stemness window (**Fig 3D-E**). Quantifying these trends among EMT status, stemness status and PD-L1 levels revealed that while hybrid E/M cells were very likely to exhibit both PD-L1 and enhanced stemness; the stemness status by itself (irrespective of EMT status) could not predict any association with PD-L1 (**Fig 3F**). The non-monotonic nature of association between EMT states and stemness was confirmed by pan-cancer data analysis of CCLE cell lines, where the stemness signature was most enriched in cells with hybrid E/M status (**Fig 3G; Fig S4E**) while no such trend was seen for a direct association of stemness with PD-L1 levels (**Fig 3H**). Together, we conclude that while hybrid E/M cells are more stem-like and immune-evasive, these two features are likely acquired independent of one another.

### Reversible resistance to anti-estrogen therapy can co-occur with an immune evasive phenotype in breast cancer

We now look more closely at the specific case of breast cancer and the connection between drug resistance and immune evasion. The emergence of reversible resistance to targeted therapy such as tamoxifen in ER+ positive breast cancer can be modulated in part by EMT-associated players such as ZEB1, miR-200 and SLUG, by virtue of their crosstalk with the estrogen receptor alpha (ERα). A cell-state switch to a hybrid E/M or a mesenchymal phenotype can enable the acquisition of a tamoxifen resistant phenotype where ERα levels are relatively low (29,30). Further, ERα has been reported to directly repress PD-L1 transcriptionally (31,32), thus raising the possibility of emergence of an immune evasion phenotype due to increased PD-L1 levels in cells that are treated with anti-estrogen therapy.

To study the emergent properties of this system, we added following components to the regulatory network in **Fig 1A** – ERα66, the major target of anti-estrogen therapy and ERα36, a mediator of anti-estrogen therapy resistance (**Fig 4A**). The interconnections between ERα66 and ERα36 were the taken from our previous study (29). Also, we included an inhibitory link from ERα66 to PD-L1 (31). We simulated this updated regulatory network in RACIPE and obtained pairwise correlations of simulated expression levels for all nodes in the network, which revealed two distinct and mutually antagonistic “teams” (33). This presumably arises due to the fact that interactions of the core EMT circuit with the estrogen response module do not exhibit any frustration (34). One team comprised EMT-inducers ZEB1 and SLUG, ERα36 and PD-L1 which were all highly correlated with one another and negatively with members of the other team which consisted of CDH1, miR-200 and ERα66 (**Fig 4B**). This pattern was consistently seen in all observed steady state solutions plotted as a heatmap, i.e., two predominant cell-states – one in which ZEB1, SLUG, ERα36, PD-L1 are high and CDH1, miR-200, ERα66 are low and *vice versa*, with a small proportion of mixed states (**Fig S5A**).

**Fig 4.**
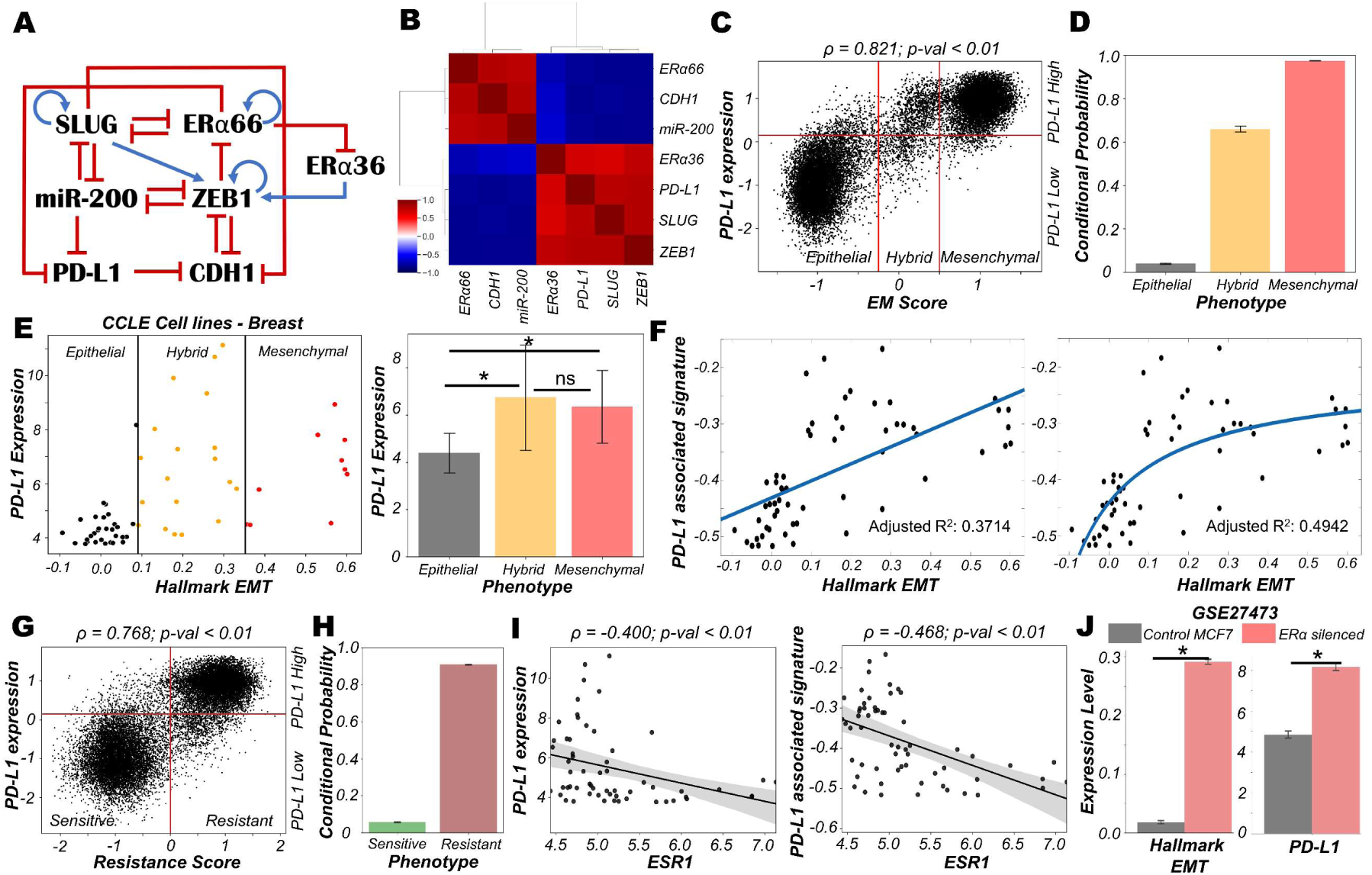
Association of high PD-L1 levels with acquisition of a reversible drug resistant phenotype in ER+ Breast cancer. **A)** Regulatory network interplay of EMT regulators, estrogen receptor isoforms (ERα66, ERα36) coupled with PD-L1. Blue arrows stand for activation links, red hammers for inhibitory links. **B)** Pairwise correlation matrix using Spearman correlations showing the existence of 2 “teams” of players – SLUG, ZEB1, ERα36, PD-L1 and CDH1, miR-200 and ERα66 – with mutually antagonistic associations. **C)** Scatter plot of PD-L1 levels and EM score. Spearman’s correlation coefficient (ρ) and corresponding p-value (p-val) are reported. **D)** Bar plot representing conditional probability of a phenotype being PD-L1 high given that it belongs to a given EMT phenotype. **E)** Scatter plot showing correlation between PD-L1 levels and the Hallmark EMT signature in breast cancer specific cell lines from CCLE. The boundaries between epithelial, hybrid and mesenchymal phenotypes are based on trisection of the entire range of Hallmark EMT scores of all cell lines in CCLE (left). Quantification of PD-L1 levels of breast cancer cell lines belonging to different EM status (right). * denotes p-val < 0.05 as assessed by a two-tailed Students t-test assuming unequal variances. **F)** Scatter plot showing linear vs Michaelis-Menten curve fit to a scatter plot of PD-L1 associated and Hallmark EMT signatures. **G)** Scatter plot of PD-L1 levels and Resistance score classified as high (>0) vs low (<0). Spearman’s correlation coefficient (ρ) and corresponding p-value (p-val) are reported. **H)** Bar plot representing conditional probability of a phenotype being PD-L1 high given that it is sensitive vs. resistant state. **I)** Scatter plot showing a significant negative correlation between PD-L1 levels and PD-L1 associated signature and ESR1 expression levels in breast cancer cell lines from CCLE. **J)** Activity/Expression levels of Hallmark EMT and PD-L1 levels in MCF7 ER+ breast cancer cells with control and ERα silenced cases (GSE27473). * denotes a statistically significant difference (p-val < 0.05) between the represented groups assessed by a two-tailed Students t-test assuming unequal variances.

To quantify the resultant phenotypes, we computed the EM score as before, and a (tamoxifen) Resistance score (= ERα36 – ERα66) (29). The resultant distribution of EM Score was visibly trimodal while that of Resistance score and PD-L1 levels were bimodal (**Fig S5B**). Plotting the EM score and Resistance scores together identified 6 phenotypes – ES (Epithelial-Sensitive), ER (Epithelial-Resistant), HS (Hybrid-Sensitive), HR (Hybrid-Resistant), MS (Mesenchymal-Sensitive), and MR (Mesenchymal-Resistant), with ES and MR being most predominant and showed a strong positive correlation between these scores (**Fig S5C**). The association between EM score and PD-L1 levels were also similarly distributed and positively correlated (**Fig 4C**) as seen earlier (**Fig 1C**). Further, a hybrid E/M or mesenchymal phenotype was more likely to be PD-L1+ as well as tamoxifen-resistant as compared to an epithelial one (**Fig 4D, Fig S5D**), consistent with previously observed trends (**Fig 1D**). These results indicate the association of properties of individual/pairs of nodes in a network remains strongly conserved, even with the addition of an extra (i.e., tamoxifen resistance) module.

Next, to validate these predictions from our simulations, we analysed breast cancer cell lines in the CCLE dataset. We designated the cell lines as epithelial, hybrid or mesenchymal by trisecting the overall range of activity of the Hallmark EMT gene set (35) (**Fig 4E** – left panel). In accordance with our model predictions, we observed PD-L1 levels in hybrid cell lines to be significantly higher than those in epithelial ones, but comparable to those in mesenchymal ones (**Fig 4E** – right panel). A similar trend was seen for enrichment of PD-L1 associated gene set (see Methods) (**Fig S5E**). This observation indicated the possibility that hybrid E/M breast cancer cell lines can be as immune evasive as mesenchymal cell lines. In other words, change in levels of PD-L1 is likely to occur in the earlier stages of EMT as compared to later stages. To further strengthen this hypothesis, we fitted a straight line (ax + b) and a Michaelis-Menten kind of a curve (ax/(b+x) + c) to a scatterplot between the ssGSEA scores of Hallmark EMT gene set and PD-L1 associated gene set (**Fig 4F**). A much better fit was observed in the latter case (adjusted R^2^ = 0.49) vs than a simple straight line fit (adjusted R^2^ = 0.37) (**Fig 4F**), indicating towards the possibility of a non-linear and saturating model of association between changes of cell phenotype along the EMT axis and consequent PD-L1 levels.

Having validated our simulation-based observations in breast cancer cell lines for EMT/PD-L1 association, we next investigated the axes of reversible drug resistance to anti-estrogen therapy and PD-L1 expression. Our simulations suggested a strong positive correlation between PD-L1 levels and resistance score (**Fig 4G**), such that resistant cells are largely PD-L1+ while sensitive ones are largely PD-L1- (**Fig 4H**). This prediction was validated in CCLE group of breast cancer cell lines, where both the expression levels of PD-L1 and corresponding ssGSEA scores of PD-L1 associated gene set were significantly negatively correlated to *ESR1* expression levels (**Fig 4I**). To gain a stronger causal evidence in support of our simulation results, we looked into a specific experimental dataset in which ERα was silenced in an ER+ breast cancer cell line, namely MCF7. We found that as ERα silencing led to a concurrent increase in the activity of the Hallmark EMT gene set as well as in PD-L1 levels (**Fig 4J**). This trend indicates that as the levels of ERα and consequently the ERα signalling activity falls, a coordinated increase in the mesenchymal nature of cells and PD-L1 levels ensues, thus leading to the resultant phenotype being more immune evasive.

Finally, we examined whether the association between PD-L1 levels, EMT and ERα levels seen *in vitro* and *in silico* holds true in patients with different subtypes of breast cancer. Analysing the TCGA cohort of patients, we found that PD-L1 and EMT signatures were positively correlated prominently in the ER+ subtype (luminal A and B) but not in ER-ones (Basal or HER2+) (**Fig S5F**). Put together, due to the interconnections among EMT status, prevalence of estrogen signalling and PD-L1, the emergence of reversible drug resistance in ER+ positive breast cancer is likely to lead to higher levels of PD-L1, thus enabling cross-resistance (i.e., tamoxifen-resistant cells being immune evasive) that further promotes their survival.

### Immune evasive properties of hybrid E/M phenotypes depends on transition trajectories in a two-dimensional (2D) EMT plane

To strengthen the association of high PD-L1 levels with hybrid E/M status of cells in other cancers, we probed the CCLE group of cell lines for lung cancer. We found that, as in the case of breast cancer, hybrid E/M cell lines exhibit significantly higher levels of PD-L1 than epithelial ones, but there was not a significant difference between hybrid E/M and mesenchymal cell lines (**Fig 5A**). Previous experimental and computational efforts has suggested SLUG+CDH1+ cells would display a hybrid E/M phenotype, especially in breast cancer (9,10). We observed this SLUG+CDH1+ profile in circulating tumour cells (CTCs) obtained from PC3 (a prostate cancer cell line, usually of a more mesenchymal status (36) (**Fig 5B**; GSE106363). This change in cell phenotype (CTC vs. cell lines) to a more hybrid E/M phenotype is accompanied by a significant increase in PD-L1 levels and concomitant decrease in ZEB1 levels (**Fig 5B**), further strengthening the association of a hybrid EM phenotype with high PD-L1 levels. Further, we analysed a publicly available expression dataset consisting of single and pooled cell prostate cancer cells along with single disseminated tumour cells. We observed a positive correlation between the Hallmark EMT program and PD-L1 associated gene set (**Fig 5C**) thus supporting our observations at a single-cell level too.

**Fig 5.**
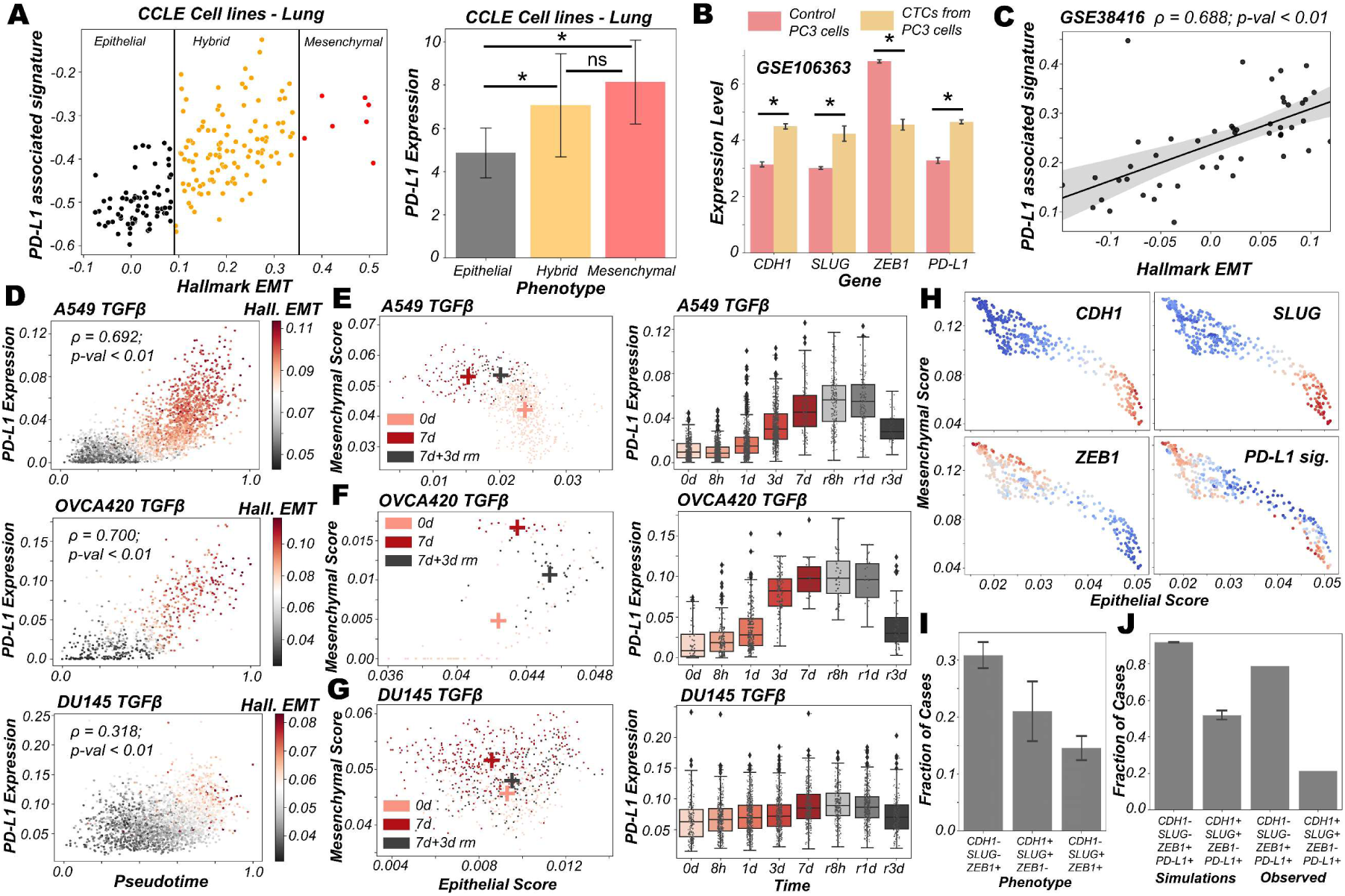
PD-L1 levels depend on the extent and direction of transition of hybrid E/M cells on 2D EM plane. **A)** Scatter plot showing PD-L1 associated and Hallmark EMT signature in lung cancer specific cell lines from CCLE. Boundaries (vertical lines) drawn are based on trisection of the entire range of Hallmark EMT scores of all cell lines in CCLE (left). Quantification of PD-L1 levels of breast cancer cell lines belonging to different EMT status (right). **B)** Expression levels of ZEB1, SLUG, CDH1 and PD-L1 in control PC3 cell line and PC3 derived CTCs (GSE106363). For A, B); * denotes a statistically significant difference (p-val < 0.05) between the represented groups assessed by a two-tailed Students t-test assuming unequal variances. **C)** Scatter plot of PD-L1 associated and Hallmark EMT signature showing a significant positive correlation in single and pooled prostate cancer cells and single DTC from metastatic prostate cancer patients (GSE38416). **D)** Scatter plot of imputed PD-L1 expression with pseudotime coloured by Hallmark EMT scores calculated on imputed gene expression data in TGFβ induced EMT in A549, OVCA420 and DU145 cell lines (GSE147405). **E-G)** 2D EM plots (left panels) showing cells of 3 different time points (day 0, day 7 and day 3 after TGFβ removal) for 3 different cell lines. + sign indicates the average epithelial and mesenchymal scores of cells belonging to the corresponding time point. Imputed PD-L1 levels over all time points plotted as boxplots (right panels). **E)** A549 **F)** OVCA420 **G)** DU145. **H)** 2D EM plots of cells from skin squamous cell carcinoma after imputation coloured by CDH1, SLUG, ZEB1 and PD-L1 associated signature. Red represents high expression while blue represents low expression (GSE110357). **I)** Abundance of top phenotypes in hybrid EM compartment shown in **Fig 1B** (simulation results). **J)** Bar plots showing the fraction of different hybrid phenotypes as measured from simulations and seen in experimental data.

Next, we delved into recent single-cell RNA-seq data that captures hybrid E/M phenotypes (37,38). We applied imputation algorithms that have been applied to EMT (39) in order to estimate PD-L1 levels in these datasets, and used the imputed levels and/or activity scores for PD-L1 associated genes. We first analysed the data collected for various cell lines *in vitro* by TGFβ treatment and withdrawal (EMT followed by MET) (37). We observed a strong correlation of PD-L1 levels with pseudo-time. This is, in turn, highly correlated with Hallmark EMT signature, as shown across different cell lines (A549, OVCA420 and DU145) from different cancer types (lung, ovarian and prostate respectively) (**Fig 5D**). Furthermore, the induction of EMT upon treatment was visibly more robust in the case of A549 and OVCA420 in comparison to DU145 cells. This was, in turn, reflected in a weaker correlation of PD-L1 levels with pseudo-time for DU145 cells as compared to other two cases (**Fig 5D**).

To get more robust information about the EMT status of the cells, we plotted cells from control case (day 0), most EMT-like state (day 7 of TGFβ treatment) and subsequent the most MET-like state (3 days after removal of TGFβ post treatment) on a two-dimensional epithelial/mesenchymal plane (**Fig 5E-G**). This 2D representation allows for deconvoluting and quantifying the activity of epithelial and mesenchymal nature of cells separately, i.e., one can independently monitor a gain of mesenchymal state and loss of epithelial nature during EMT. On this 2D plane, A549 cells showed both a loss of epithelial traits and a gain of mesenchymal ones as they were treated with TGFβ (**Fig 5E;** left). It is important to note here that the 2D plane only captures relative changes in epithelial and mesenchymal nature; without an external reference point, it is difficult to ascertain the absolute EMT status of cells at a given timepoint. Furthermore, upon MET (TGFβ withdrawal), A549 cells partly regained their epithelial nature, without any discernible change in mesenchymal nature. Thus, these cells can be referred to as the hybrid E/M state with respect to the untreated case. Consistently, PD-L1 levels increased as cells progressed through EMT (TGFβ treatment time-points) and decreased as they began their return (TGFβ withdrawal post-treatment). PD-L1 levels in cells 3 days after TGFβ withdrawal (i.e., hybrid E/M) were still higher as compared to untreated cells (**Fig 5E**; right). This suggests that cells fine-tune their PD-L1 levels as they proceed through EMT/MET, thereby enabling hybrid E/M cells to be sufficiently immune-evasive. More interesting trends were seen for the trajectory of OVCA420 cells on the 2D EMT plane (**Fig 5F**) which did not lose any epithelial characteristics upon EMT induction, but gained mesenchymal nature, making the resultant phenotype at day 7 post-treatment a largely hybrid E/M population (**Fig 5F**; left). PD-L1 levels robustly increased, bolstering further evidence for hybrid E/M phenotypes having higher levels of PD-L1 as compared to their more epithelial counterparts (**Fig 5F**; right). As compared to A549 and OVCA420, DU145 cells showcased a much weaker induction of EMT. Consequently, no strong increase in the levels of PD-L1 was observed (**Fig 5G**). This analysis of three different cell lines at multiple time points at an individual-cell level strongly supports our predictions of increased PD-L1 levels being a hallmark of hybrid E/M phenotypes.

Having shown that PD-L1 levels observed in hybrid E/M cells depends on the extent and direction of transitions on a 2D EM plane (**Fig 5G**), we proceeded to interrogate possible mechanistic basis of heterogeneous hybrid E/M phenotypes using data recently reported *in vivo* in squamous cell cancer (38). We applied imputation algorithm MAGIC (39) on the data, we plotted it on 2D EMT plane, and observed an expected negative correlation between the epithelial and mesenchymal programs (**Fig 5H**). A large proportion of cells seen here were CDH1+SLUG+ZEB1- and ZEB1+SLUG-CDH1-, which can be interpreted as two different versions of hybrid E/M phenotypes – the former being relatively more epithelial (“early hybrid”) and the latter being more pushed towards a mesenchymal end (“late hybrid”). Interestingly, CDH1+SLUG-ZEB1- (“strongly epithelial”) and CDH1-SLUG+ZEB1+ (“strongly mesenchymal”) phenotypes were seen only sporadically in this dataset. Among the two hybrid E/M phenotypes, PD-L1 associated gene signature was enriched in a substantial proportion of cells, with a larger number of PD-L1+ cases in the ZEB1+SLUG-CDH1-hybrid subpopulation (**Fig 5H**), thereby identifying context-specific scenarios for heterogeneous hybrid E/M subsets which may manifest immune-evasive properties.

We attempted to offer a mechanistic reason for enrichment of PD-L1+ cases in the ZEB1+SLUG-CDH1-population using our network-based simulations. When we classified all of the steady state solutions by binarizing and considering only the ZEB1, SLUG and CDH1 status in hybrid EM compartment shown in **Fig 1C**, we found that most of the solutions mapped onto a ZEB1+SLUG-CDH1-phenotype closely followed by CDH1+SLUG+ZEB1-phenotype (**Fig 5I**). Furthermore, our parameter-agnostic modelling framework was able to recapitulate the qualitative association of ZEB1+SLUG-CDH1-hybrid E/M phenotype being more likely to be PD-L1+ than the CDH1+ SLUG+ZEB1-hybrid E/M phenotype (**Fig 5J**). While further rigorous experiments are required to substantiate this observation about heterogeneity in hybrid E/M subpopulations in terms of their PD-L1 levels, our analysis elucidates how deciphering the emergent dynamics of the small-scale regulatory networks can explain reported heterogeneity in hybrid E/M state at a single-cell level.

## Discussion

Cellular processes are controlled by numerous regulatory feedback loops and mechanisms that maintain a dynamic equilibrium, thus enabling cells to adapt to various internal and external fluctuations. The expression of PD-L1 on cell surface is one such mechanism, that keeps inflammatory responses from uncontrolled activation by providing necessary brakes (2). Tumour cells exploit this checkpoint to escape from both immunological detection and elimination. PD-L1 on cancer cells’ surface enables them to inhibit T-cell activation, while simultaneously causing them to be exhausted, eventually preventing cancer cells from being targeted by activated T cells (2). High PD-L1 has been exhibited in circulating tumor cells as well across cancers (40,41), and EMT has been associated with higher PD-L1 levels (5). Here, we have investigated PD-L1 levels in hybrid E/M phenotypes, given their higher fitness for metastasis and evasion of various treatment options.

Through *in silico* simulations for underlying networks incorporating crosstalk between PD-L1 and EMT regulators, we observed that hybrid E/M states can show high PD-L1 levels similar to those seen in ‘full EMT’ (mesenchymal) phenotype. This model prediction is substantiated by analysis of gene expression pan-cancer datasets both at individual and bulk RNA-seq levels. We further show that EMT/MET can alter PD-L1 status reversibly in cancer cells, a trend validated in multiple *in vitro* datasets. Intriguingly, while hybrid E/M phenotype was found to be associated with both enhanced PD-L1 and higher stemness, a direct association between PD-L1 levels and stemness was not found. In contrast, residual drug-resistant cells that survive treatment with anti-estrogen therapy in breast cancer are likely to also harbour a hybrid E/M phenotype and higher levels of PD-L1. Our results thus highlight another dimension of the high metastatic fitness of hybrid E/M cells: their immune-suppressive traits.

We acknowledge, as with all models, the limitations of our analysis. We considered here a minimalistic regulatory network that captures key regulatory nodes of interest, and thus is far from being comprehensive. On the one hand, our model recapitulates key observations especially including previously reported associations between EMT and PD-L1 levels; on the other, it provides testable predictions, regarding hybrid E/M states being likely to be PD-L1 positive. The overarching positive correlations between EMT and PD-L1 levels across a majority of the cancers in TCGA shows the broad applicability of our conclusions; also, these were supported in by the analysis of the CCLE and of specific datasets dealing with perturbations. However, our generic model cannot explain some specific exceptions, specifically why certain cancer types such as TGCT, PCPG, SARC and SKCM do not show strong correlation of PD-L1 with either epithelial or mesenchymal programmes. Interestingly, some cancers of mesenchymal origins (LGG, GBM) show positive correlations of PD-L1 with epithelial signatures, suggesting the association of a hybrid E/M state with maximal PD-L1 levels. Finally, our extended model, obtained by considering the additional players OCT4, LIN28, miR-145 and let-7, finds no significant association between stemness and PD-L1 levels. Various additional nodes in the network not considered here may alter this trend, which may then explain previously reported correlations between immune evasion and stemness (42). However, our model predicted an overlap between immune evasion and resistance to targeted therapy in breast cancer. Thus, future efforts are needed to understand these context-specific scenarios in terms of interplay between hybrid E/M phenotypes, stemness, targeted therapy resistance and immune evasion.

There has been a recent surge of interest in hybrid E/M phenotypes, and their precise identification is an active area of research, thus calling for more rigorous, preferably quantitative, definition(s). Such definitions, instead of purely descriptive characterization of hybrid E/M cells, can offer new conceptual insights into markers and features of *bona fide* hybrid E/M cells (43). Here, we have used two quantitative strategies based on bulk and single-cell RNA-seq data to demarcate hybrid E/M cells from epithelial and mesenchymal ones. First, we used a two-dimensional metric to quantify EMT (44): epithelial and mesenchymal axes, so that individual changes along those both axes can be deconvoluted and different paths to EMT/MET can be seen, which are beyond the scope for existing scoring metrics for EMT transcriptomes (45). This strategy facilitated us to map changes in PD-L1 levels in cells as a function of their trajectory on the 2D plane and strengthened the association between different possible hybrid E/M state(s) and enhanced PD-L1 levels. Second, we defined more absolute boundaries for EMT enrichment using a cohort of CCLE cell lines to classify them into epithelial, hybrid and mesenchymal categories. Such approaches allow us to also characterise underlying manifestations and reasons for heterogeneity within the hybrid E/M phenotypes.

## Materials and methods

### RACIPE simulations on gene regulatory networks

RACIPE simulations were performed on gene regulatory networks shown in **Fig 1A, FigS4C** and **Fig 4A**. The gene regulatory links for these networks have been listed in **Table S3**. We generated steady state gene expression values for 100 initial conditions each for 10,000 parameter sets using RACIPE. Euler’s Method was used to numerically integrate the coupled differential equations representing the various circuits. RACIPE takes a topology file as an input and samples the parameters for dynamical simulations from a biologically relevant ranges (**Table S4**). Depending on the exact parameter set, a single model has the potential to give rise to one or more stable steady-state solutions, for a given set of initial conditions. For all further analysis, we have considered up to 5 stable steady-state solutions as majority of the parameter sets results in <5 steady state solutions. RACIPE generates the steady-state values in the log2 scale, which we have further converted into z-scores by using the equation represented in **Equation 1** for a better comparative study of the expression of each gene node.

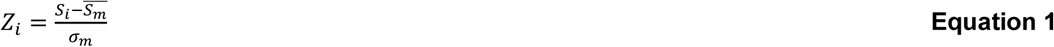

where, *Z*_*i*_ = Z normalised expression value, S_i_ = each steady state value of a given node, 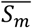 = combined mean of untransformed expression values and *σ*_m_ = combined standard deviation of untransformed expression values.

PCA was performed on all the z-normalised steady-state solutions. We identified 3 optimal clusters by performing hierarchical clustering on the z-normalised RACIPE data. Contributions of the various node to the principal component axes PC-1 and PC-2 presented in **Table S1**.

For over-expression or down-expression of specific genes as shown in **Fig 2D, E**, we simulated and normalised the gene regulatory network as shown in **Fig 1A** as above with the added specification to either over-express or down-express the node of interest (miR-200 or SLUG) by 20-fold.

### Gene regulatory network for crosstalk between EMP and PD-L1 and EM score calculation

The scores were calculated by difference in normalized expression values of node representing mesenchymal (M) and epithelial (E) signatures. EM score = (ZEB1 + SLUG - miR200 - CDH1)/4. Subsequently based on the minima values in the distribution, steady state solutions were categorised into epithelial (<-0.25), hybrid E/M (−0.25 to 0.5), and mesenchymal (>0.5) represented in **Fig 1B**. The same metric was used to calculate EM score for calculation of EM scores for gene regulatory networks shown in **Fig S4C** and **Fig 4A**. Boundaries for categorisation were kept constant for these circuits too. Steady state solutions having PD-L1 levels > 0 were termed as PD-L1+ and those with < 0 were termed as PD-L1-.

### Gene regulatory network for crosstalk between stemness circuit, EMP and PD-L1 and stemness score calculation

We have considered a gene regulatory network shown in **Fig S4C** in which 5 nodes of our core regulatory network are present along with 4 other nodes (OCT4, miR-145, LIN28, and let7) which represents the key players of stemness signature (**Table S1)**. The stemness scores (SN) were calculated by difference in normalized expression values of node representing stem-like and non-stem-like signatures: (LIN28 + OCT4 – let7 – miR145)/4. Subsequently based on the minima values in the distribution, steady state solutions were categorised into non-stem-like (SN score < −0.5 and SN score > 0.5), and stem-like (SN score = −0.5 to 0.5) represented in **Fig 3C**. Steady state solutions having PD-L1 levels > 0 were termed as PD-L1+ and those with < 0 were termed as PD-L1-. EM scores and EM status classification thresholds were kept the same as for the core gene regulatory network (**Fig 1A**).

### Gene regulatory network for therapy resistance in breast cancer and Resistance score calculation

We have considered a gene regulatory network shown in **Fig 4A** in which 5 nodes of our core regulatory network are present along with 2 other nodes (ERɑ66 and ERɑ36) which represents the key players proxy of tamoxifen sensitivity and resistance respectively (**Table S1**). The resistance score was calculated as (ERɑ36 – ERɑ66)/2. Subsequently based on the distribution, cells were categorised into sensitive (resistance score < 0), and resistance (resistance score > 0). As the minima for PD-L1 expression was at 0.15 instead of 0, for all the analysis with respect to **Fig 4A**, PD-L1 expression above 0.15 was considered as PD-L1+ and those < 0.15 as PD-L1 negative. EM scores and EM status classification thresholds were kept the same as for the core gene regulatory network (**Fig 1A**).

### Stochastic simulations and Landscape construction

We simulated the gene regulatory network in **Fig 1A** using the Euler-Maruyama method for a representative parameter set (**Table S5**) that showed the co-existence of 3 phenotypes: epithelial with low levels of PD-L1; hybrid E/M with high levels of PD-L1 and mesenchymal with high levels of PD-L1. The corresponding equation is as follows:

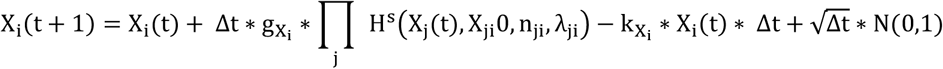

The equation is just a discrete form of the ODE presented before, with an addition of the noise term 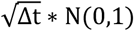, where Δt is the time step and N(0,1) is a normal random variable with mean 0 and standard deviation 1. For the parameter set, we simulated the network for 100 different initial conditions sampled uniformly from the range 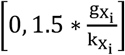. We then normalized the trajectories using the mean and standard deviation of each node expression obtained from RACIPE and converted the trajectories to EM scores and PD-L1 levels in order to classify them into the observed phenotypes.

Using these trajectories, we constructed obtained a probability density (P) of the EM-PD-L1 score pairs and constructed a potential landscape by calculating the pseudo potential as – log (P) (46).

### sRACIPE simulations

sRACIPE simulations were performed on the gene regulatory network in **Fig 1A** with a fixed amount of noise. We have used the webserver of Gene Circuit Explorer (GeneEx) to simulate stochastic time evolution dynamics of our core gene regulatory circuits (22). Parameter values used for the simulation are presented in **Table S6**.

### Gene expression data analysis of bulk microarray / RNA-seq data

Publicly available microarray and RNA-Seq datasets were obtained from GEO. Single-sample gene set enrichment analysis (ssGSEA) (47) was performed on the Hallmark signalling pathways gene signatures from MSigDB (Molecular Signatures Database) (35) to estimate the activity of pathway.

### PD-L1 associated signature(s)

To generate PD-L1 associated signatures, we selected top correlated genes (Spearman correlation > 0.5; p-val < 0.01) with PD-L1 levels across at least 15 out of the 27 cancer types considered for the study (excluding TGCT, PCPG, SARC, SKCM and KIRC as these did not show any consistent association with either the epithelial or mesenchymal metrics). The criteria of 15 out of 27 was relaxed to 13 out of 27 for analysis of **Fig 5H-J** as the signature from the more stringent signature did not have sufficient power to distinguish the cell populations.

### scRNA Seq data analysis

Read counts for scRNA seq datasets were downloaded from GEO datasets and imputation was performed on these datasets using MAGIC algorithm (39). Computation of activity scores of signature gene sets on the imputed gene expression matrices were done using AUCell (48). For computation of epithelial and mesenchymal scores tumour specific gene lists were used from the KS EMT scoring metric (45).

## Acknowledgement

MKJ was supported by Ramanujan Fellowship (SB/S2/RJN-049/2018) awarded by Science and Engineering Research Board (SERB), Department of Science & Technology, Government of India. KH acknowledges support from Prime Ministers’ Research Fellowship (PMRF). SS acknowledges support by KVPY fellowship.

## Author contributions

SS, SPN, KH, PP, AK, SM performed research; SS and SPN prepared first draft of the manuscript; all authors analysed data and edited the manuscript. MKJ and HL designed research; MKJ supervised research.

## Supplemental figures

**Fig S1.**
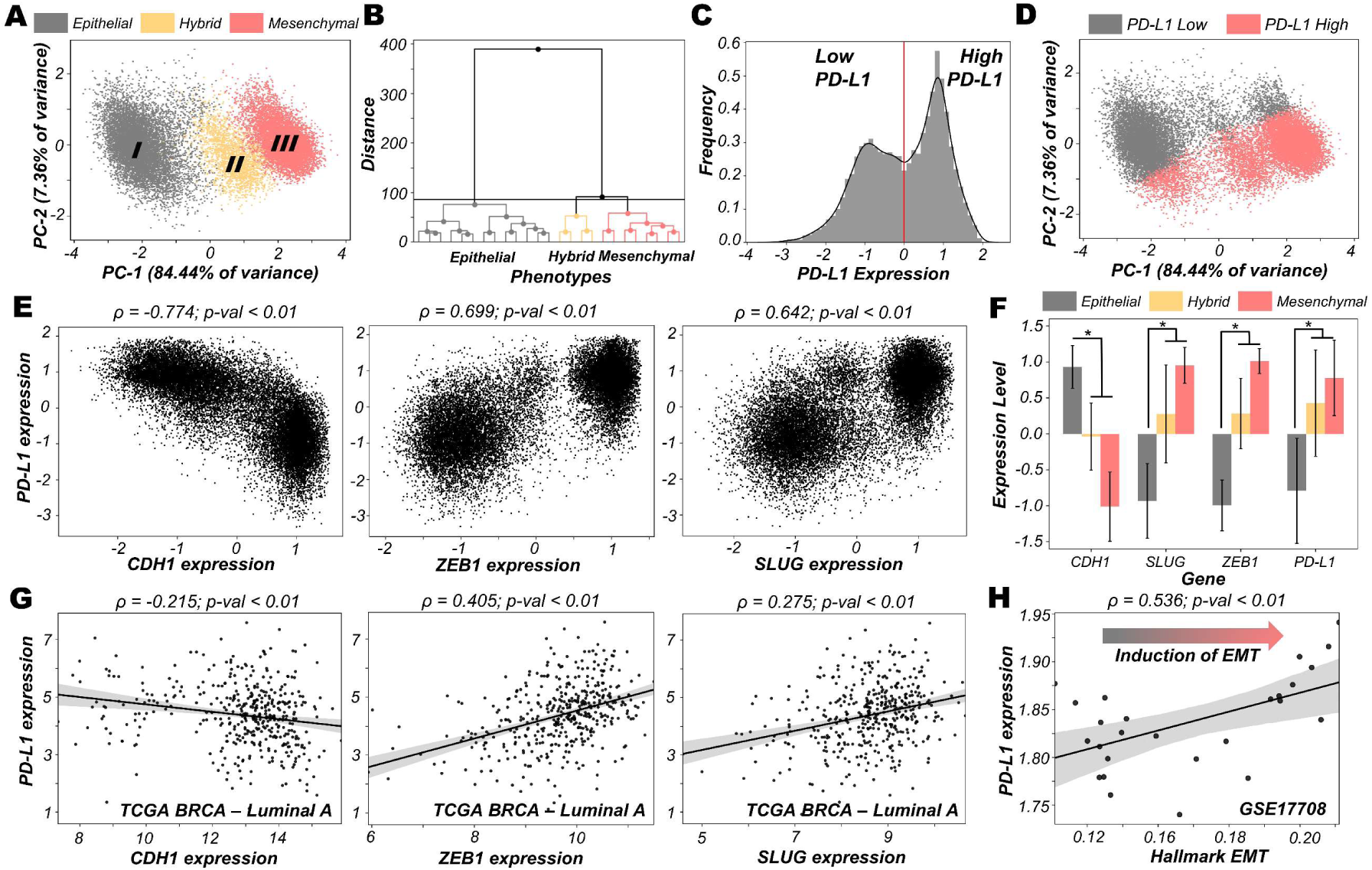
EMT regulatory network coupled with PD-L1. **A)** PCA (Principal Component Analysis) plot showing the presence of different clusters emerging from z-normalised scores from RACIPE analysis. Composition of PC1 and PC2 are listed in Table S3. **B)** Hierarchical clustering for z normalised RACIPE output. **C)** Density histogram of PD-L1 expression fitted with kernel density estimate, showing bimodality. Red lines show the partition between PD-L1 high and PD-L1 low. **D)** PCA plot coloured by PD-L1 high vs. PD-L1 low levels, showing the enrichment of high PD-L1 levels in hybrid E/M and mesenchymal phenotypes, and that of low PD-L1 levels in an epithelial phenotype. **E)** Simulation results showing scatter plot of PD-L1 expression with CDH1, ZEB1, and SLUG, as obtained from RACIPE simulations. Spearman’s correlation coefficient (ρ) and corresponding p-value (p-val) are reported. **F)** Bar graph showing expression of CDH1, SLUG, ZEB1 and PD-L1 in corresponding phenotypes (defined based on EM scores) respectively. **G)** Scatter plot showing experimental validation from TCGA BRCA – Luminal A cohort of patients of correlations between expression of PD-L1 with CDH1, ZEB1, and SLUG, which was earlier represented in E). **H)** Scatter plot showing positive correlation between PD-L1 expression and Hallmark EMT signature in A549 lung adenocarcinoma cells treated with TGFβ to induce EMT over 3 days (GSE17708). * denotes a statistically significant difference (p-val < 0.05) between the represented groups assessed by a two-tailed Students t-test assuming unequal variances.

**Fig S2.**
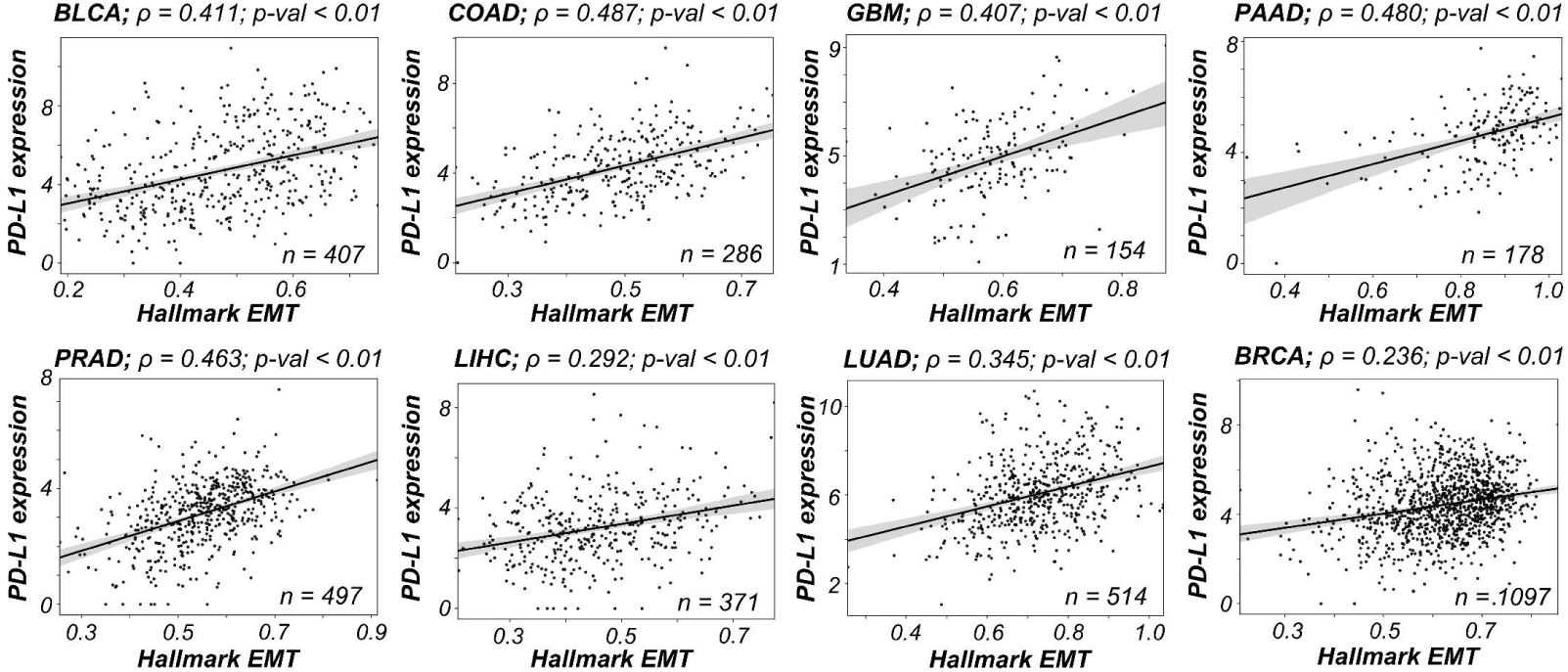
Clinical evidence supporting mathematical model predictions. Scatter plots between expression levels of PD-L1 and Hallmark EMT in representative TCGA cancer types. Spearman’s correlation coefficient (ρ) and corresponding p-value (p-val) are reported.

**Fig S3.**
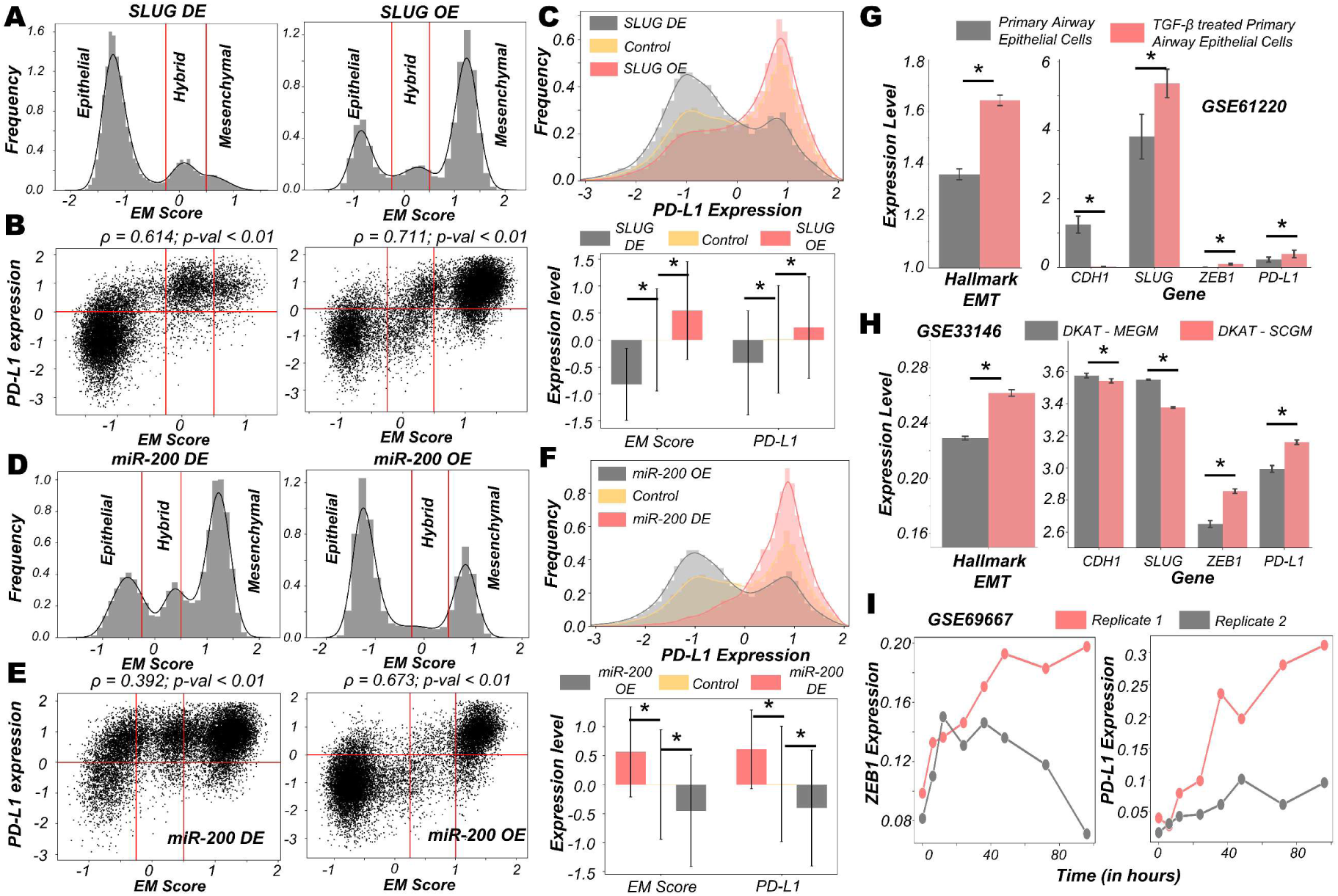
Dynamics upon perturbation of core regulatory network. **A-C)** Upon miR-200 down expression (DE) and miR-200 over expression (OE): **A)** density histogram of EM Score fitted with kernel density estimate; **B)** Scatter plot of PD-L1 expression and EM score; **C)** Density histogram of PD-L1 expression fitted with kernel density estimate and Bar graph showing change in expression of EM score and PD-L1. **D-F)** Same as A-C but for SLUG DE and SLUG OE. Horizontal red line shows the partition between PD-L1 expression level being high and low. Vertical red lines show a partition between phenotypes: Epithelial, Hybrid E/M, and Mesenchymal based on EM score. Spearman’s correlation coefficient (ρ) and corresponding p-value (p-val) are given. **G)** Activity/Expression levels of Hallmark EMT and PD-L1 levels in non-cancerous airway epithelial cells where EMT has been induced (GSE61220). **H)** Activity/Expression levels of Hallmark EMT and PD-L1 levels in triple negative breast cancer (DKAT) cells grown in either epithelial growth medium (MEGM) or stromal growth medium (SCGM) (GSE33146). **I)** Expression levels of ZEB1 and PD-L1 in A549 lung cancer cells with EMT induced via TGFβ (GSE27473). * denotes a statistically significant difference (p-val < 0.05) between the represented groups assessed by a two-tailed Students t-test assuming unequal variances.

**Fig S4.**
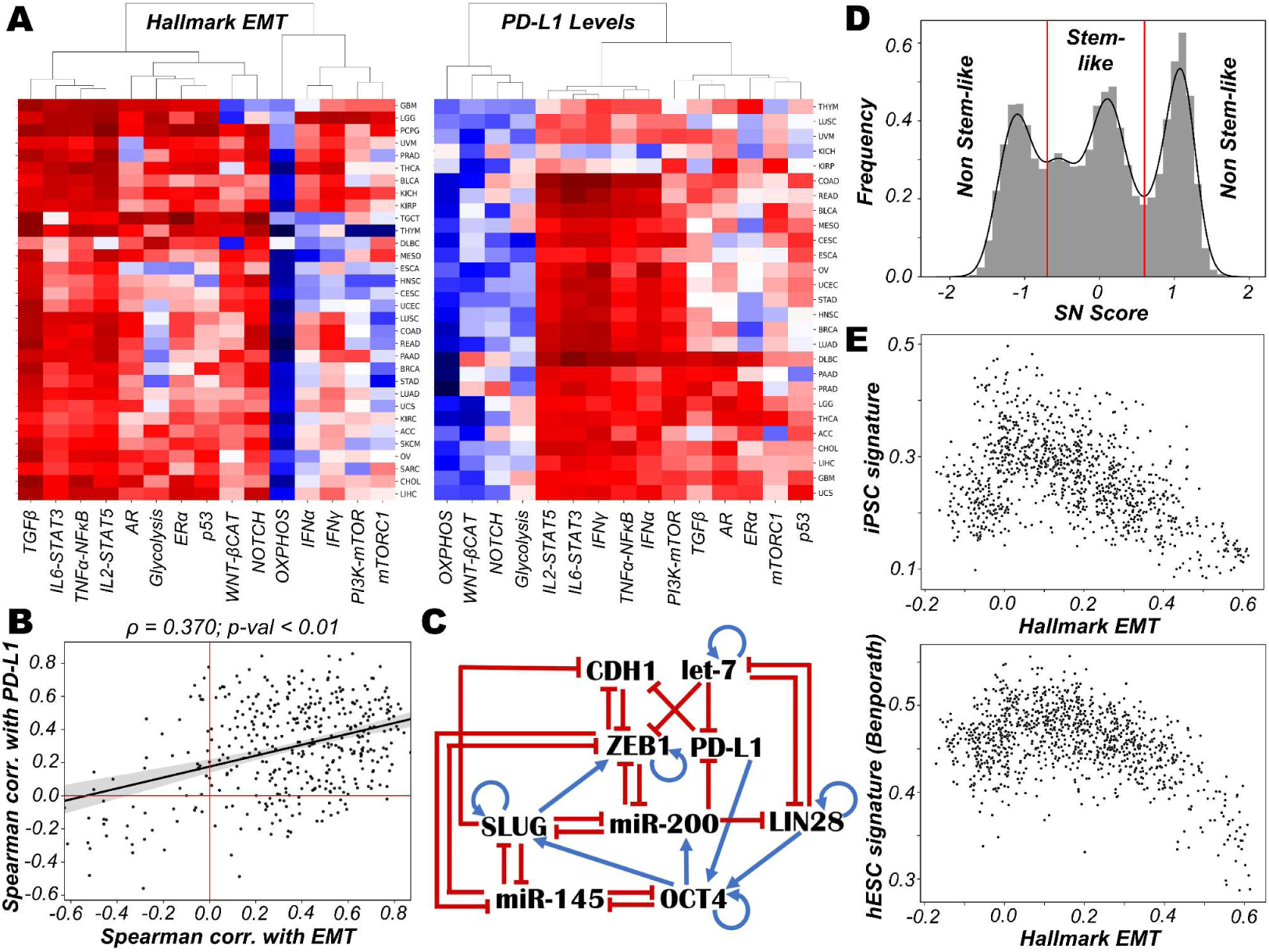
Different pathways that may influence the EMT/PD-L1 association. **A)** Heatmap showing Spearman’s correlation between various signalling pathways and Hallmark EMT/PD-L1 levels respectively. Spearman’s correlation coefficient (ρ) and e corresponding p-value (p-val) are reported. **B)** Scatter plots between the Spearman’s correlation of expression levels of PD-L1 and spearman correlation of Hallmark EMT showing the concordance between two heatmaps in (A). **C)** Schematic representation of stemness circuit diagram with nodes representing various EMT, immune evasion, and stemness signature players. **(D)** Density histogram of Stemness Score (SN score) (LIN28 + OCT4 – let7 – miR145)/4 fitted with kernel density estimate showing predominantly a trimodal distribution. Vertical red lines show the partition between stem-like and non-stem-like based on SN score, where intermediate levels of SN score lie within the ‘stemness window’. **E)** Scatter plots between expression levels of PD-L1 with iPSC signature and hESC signature (Ben-porath et al. 2008) respectively in CCLE datasets.

**Fig S5.**
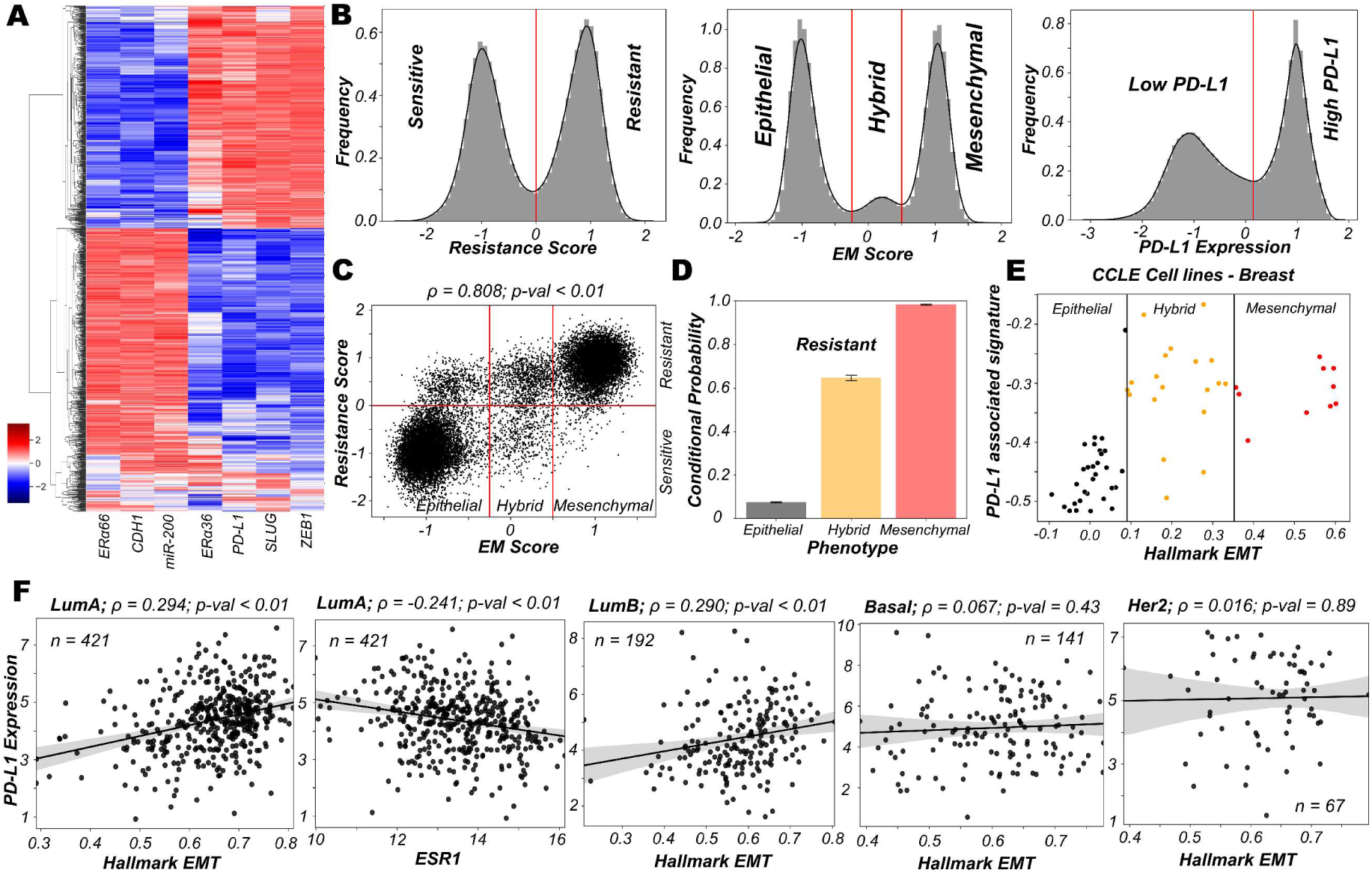
Chacterisation of the association of high PD-L1 levels upon acquisition of a reversible drug resistant phenotype in ER+ breast cancer. **A)** Heatmap showing stable steady-state solutions for the gene regulatory network shown in Fig 4A obtained via RACIPE. **B)** Frequency density histograms for Resistance score, EM score and PD-L1 levels. The red vertical lines discretise the continuous distributions to distinct phenotypes based on the minima found in the distribution. **C)** Scatter plot showing a strong association of the EM score with the resistance score followed by classification to 6 possible phenotypes. Spearman’s correlation coefficient (ρ) and corresponding p-value (p-val) are reported. **D)** Bar plot representing conditional probability of a phenotype being a resistant phenotype given that it belongs to a given EMT status. **E)** Scatter plot showing correlation between PD-L1 associated gene set and the Hallmark EMT signature in breast cancer specific cell lines from CCLE. The boundaries between epithelial, hybrid and mesenchymal phenotypes are based on trisection of the entire range of Hallmark EMT scores of all cell lines in CCLE. **F)** Scatter plots between expression levels of PD-L1 and Hallmark EMT across different subtypes in TCGA BRCA cohort of patients. The scatter plot between expression levels of PD-L1 and ESR1 has also been shown for Luminal A subtype of breast cancer. Spearman’s correlation coefficient (ρ) and corresponding p-value (p-val) are reported.

**Fig S6.**
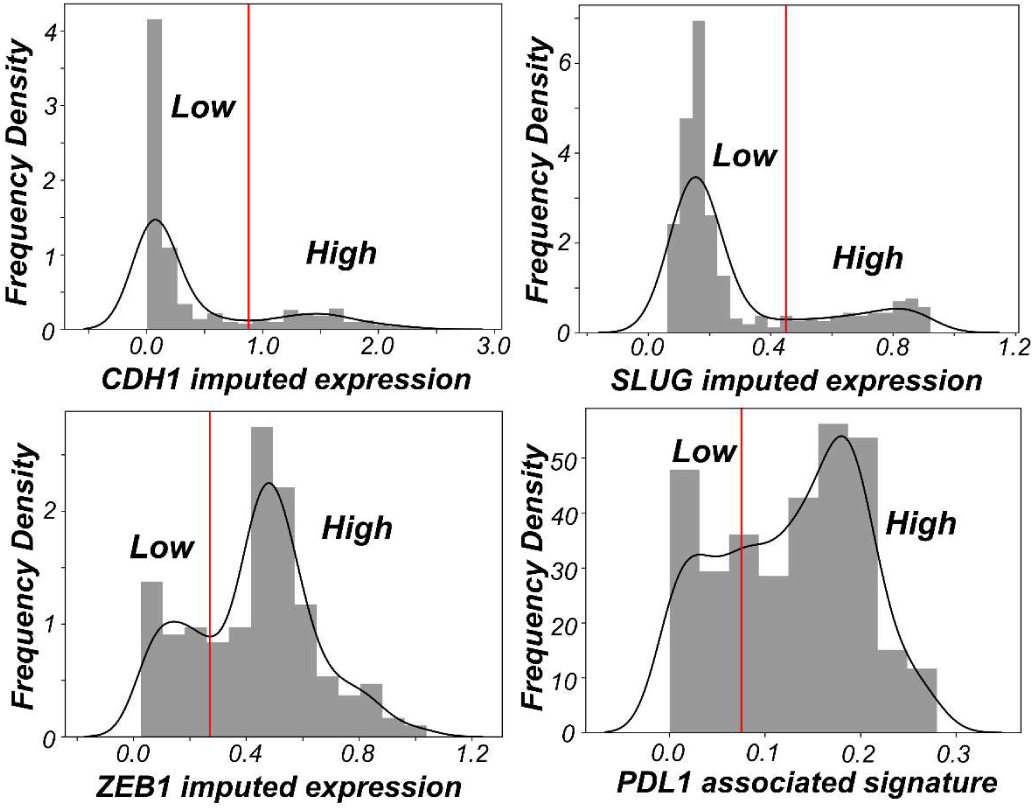
Discretisation of imputed gene expression/activity scores for CDH1, ZEB1, SLUG and PD-L1 associated signature. The cutoff of high vs low is decided based on the minima in the bimodal distributions seen; expect in the case of PD-L1 associated signature, where it is decided as the average of the two relatively shallow minimas in the distribution.

## References

1. Binnewies M, Roberts EW, Kersten K, Chan V, Fearon DF, Merad M, et al. Understanding the tumor immune microenvironment (TIME) for effective therapy. Nat Med. 2018;24:541–50.

2. Sun C, Mezzadra R, Schumacher TN. Regulation and Function of the PD-L1 Checkpoint. Immunity. 2018;48:434–52.

3. Hashimoto M, Kamphorst AO, Im SJ, Kissick HT, Pillai RN, Ramalingam SS, et al. CD8 T Cell Exhaustion in Chronic Infection and Cancer: Opportunities for Interventions. Annu Rev Med. 2018;69:301–18.

4. Chen L, Gibbons DL, Goswami S, Cortez MA, Ahn Y, Byers L a, et al. Metastasis is regulated via microRNA-200/ZEB1 axis control of tumour cell PD-L1 expression and intratumoral immunosuppression. Nat Commun. 2014;5:5241.

5. Dongre A, Rashidian M, Reinhardt F, Bagnato A, Keckesova Z, Ploegh HL, et al. Epithelial-to-mesenchymal transition contributes to immunosuppression in breast carcinomas. Cancer Res. 2017;77:3982–9.

6. Guo Y, Lu X, Chen Y, Rendon B, Mitchell RA, Cuatrecasas M, et al. Zeb1 induces immune checkpoints to form an immunosuppressive envelope around invading cancer cells. Sci Adv. 2021;7:eabd7455.

7. Jolly MK, Somarelli JA, Sheth M, Biddle A, Tripathi SC, Armstrong AJ, et al. Hybrid epithelial/mesenchymal phenotypes promote metastasis and therapy resistance across carcinomas. Pharmacol Ther. 2019;194:161–84.

8. Lu M, Jolly MK, Levine H, Onuchic JN, Ben-Jacob E. MicroRNA-based regulation of epithelial–hybrid–mesenchymal fate determination. Proc Natl Acad Sci. 2013;110:18144–9.

9. Subbalakshmi AR, Sahoo S, Biswas K, Jolly MK. A computational systems biology approach identifies SLUG as a mediator of partial Epithelial-Mesenchymal Transition (EMT). Cells Tissues Organs. 2021;in press.

10. Sterneck E, Poria DK, Balamurugan K. Slug and E-Cadherin: Stealth Accomplices? Front Mol Biosci. 2020;7:138.

11. Burk U, Schubert J, Wellner U, Schmalhofer O, Vincan E, Spaderna S, et al. A reciprocal repression between ZEB1 and members of the miR-200 family promotes EMT and invasion in cancer cells. EMBO Rep. 2008;9:582–9.

12. Wels C, Joshi S, Koefinger P, Bergler H, Schaider H. Transcriptional activation of ZEB1 by Slug leads to cooperative regulation of the epithelial-mesenchymal transition-like phenotype in melanoma. J Invest Dermatol. 2011;131:1877–85.

13. Liu YN, Yin JJ, Abou-Kheir W, Hynes PG, Casey OM, Fang L, et al. MiR-1 and miR-200 inhibit EMT via Slug-dependent and tumorigenesis via Slug-independent mechanisms. Oncogene. 2013;32:296–306.

14. Chen L, Xiong Y, Li J, Zheng X, Zhou Q, Turner A, et al. PD-L1 Expression Promotes Epithelial to Mesenchymal Transition in Human Esophageal Cancer. Cell Physiol Biochem. 2017;42:2267–80.

15. Huang B, Lu M, Jia D, Ben-Jacob E, Levine H, Onuchic JN. Interrogating the topological robustness of gene regulatory circuits by randomization. PLoS Comput Biol. 2017;13:e1005456.

16. Juneja VR, McGuire KA, Manguso RT, LaFleur MW, Collins N, Nicholas Haining W, et al. PD-L1 on tumor cells is sufficient for immune evasion in immunogenic tumors and inhibits CD8 T cell cytotoxicity. J Exp Med. 2017;214:895–904.

17. Hong T, Watanabe K, Ta CH, Villarreal-Ponce A, Nie Q, Dai X. An Ovol2-Zeb1 Mutual Inhibitory Circuit Governs Bidirectional and Multi-step Transition between Epithelial and Mesenchymal States. PLOS Comput Biol. 2015;11:e1004569.

18. George JT, Jolly MK, Xu S, Somarelli JA, Levine H. Survival outcomes in cancer patients predicted by a partial EMT gene expression scoring metric. Cancer Res. 2017;77:6415–28.

19. Hong D, Messier TL, Tye CE, Dobson JR, Fritz AJ, Sikora KR, et al. Runx1 stabilizes the mammary epithelial cell phenotype and prevents epithelial to mesenchymal transition. Oncotarget. 2017;8:17610–27.

20. Celià-Terrassa T, Meca-Cortés Ó, Mateo F, De Paz AM, Rubio N, Arnal-Estapé A, et al. Epithelial-mesenchymal transition can suppress major attributes of human epithelial tumor-initiating cells. J Clin Invest. 2012;122:1849–68.

21. Padda SK, Riess JW, Schwartz EJ, Tian L, Kohrt HE, Neal JW, et al. Diffuse high intensity PD-L1 staining in thymic epithelial tumors. J Thorac Oncol. 2015;10:500–8.

22. Kohar V, Lu M. Role of noise and parametric variation in the dynamics of gene regulatory circuits. npj Syst Biol Appl. 2018;4:40.

23. Tripathi S, Levine H, Jolly MK. The Physics of Cellular Decision-Making during Epithelial-Mesenchymal Transition. Annu Rev Biophys. 2020;49:1–18.

24. Gonzalez DM, Medici D. Signaling mechanisms of the epithelial-mesenchymal transition. 2014;7:1–17.

25. Yi M, Niu M, Xu L, Luo S, Wu K. Regulation of PD-L1 expression in the tumor microenvironment. J Hematol Oncol. 2021;14:10.

26. Hai Ping P, Feng Bo T, Li L, Nan Hui Y, Hong Z. IL-1β/NF-kb signaling promotes colorectal cancer cell growth through miR-181a/PTEN axis. Arch Biochem Biophys. 2016;604:20–6.

27. Morel A-P, Lièvre M, Thomas C, Hinkal G, Ansieau S, Puisieux A. Generation of breast cancer stem cells through epithelial-mesenchymal transition. PLoS One. 2008;3:e2888.

28. Jolly MK, Huang B, Lu M, Mani SA, Levine H, Ben-Jacob E. Towards elucidating the connection between epithelial - mesenchymal transitions and stemness. J R Soc Interface. 2014;11:20140962.

29. Sahoo S, Mishra A, Kaur H, Hari K, Muralidharan S, Mandal S, et al. A mechanistic model captures the emergence and implications of non-genetic heterogeneity and reversible drug resistance in ER+ breast cancer cells. NAR Cancer. 2021;3:zcab027.

30. Alves CL, Elias D, Lyng MB, Bak M, Ditzel HJ. SNAI2 upregulation is associated with an aggressive phenotype in fulvestrant-resistant breast cancer cells and is an indicator of poor response to endocrine therapy in estrogen receptor-positive metastatic breast cancer. Breast Cancer Res. 2018;20:60.

31. Liu L, Shen Y, Zhu X, Lv R, Li S, Zhang Z, et al. ERα is a negative regulator of PD-L1 gene transcription in breast cancer. Biochem Biophys Res Commun. 2018;505:157–61.

32. Huhn D, Marti-Rordigo P, Mouron S, Hansel CS, Tschapalda K, Porebski B, et al. Prolonged estrogen deprivation triggers an immunosuppressive phenotype in breast cancer cells. Mol Oncol. 2021;in press.

33. Chauhan L, Ram U, Hari K, Jolly MK. Topological signatures in regulatory network enable phenotypic heterogeneity in small cell lung cancer. Elife. 2021;10:e64522.

34. Tripathi S, Kessler DA, Levine H. Biological Networks Regulating Cell Fate Choice Are Minimally Frustrated. Phys Rev Lett. 2020;125:088101.

35. Liberzon A, Subramanian A, Pinchback R, Thorvaldsdóttir H, Tamayo P, Mesirov JP. Molecular signatures database (MSigDB) 3.0. Bioinformatics. 2011;27:1739–40.

36. Roca H, Hernandez J, Weidner S, McEachin RC, Fuller D, Sud S, et al. Transcription Factors OVOL1 and OVOL2 Induce the Mesenchymal to Epithelial Transition in Human Cancer. PLoS One. 2013;8:e76773.

37. Cook DP, Vanderhyden BC. Context specificity of the EMT transcriptional response. Nat Commun. 2020;11:2142.

38. Pastushenko I, Brisebarre A, Sifrim A, Fioramonti M, Revenco T, Boumahdi S, et al. Identification of the tumour transition states occurring during EMT. Nature. 2018;556:463–8.

39. van Dijk D, Sharma R, Nainys J, Yim K, Kathail P, Carr AJ, et al. Recovering Gene Interactions from Single-Cell Data Using Data Diffusion. Cell [Internet]. Elsevier Inc.; 2018;174:716-729.e27. Available from: https://doi.org/10.1016/j.cell.2018.05.061

40. Bergmann S, Coym A, Ott L, Soave A, Rink M, Janning M, et al. Evaluation of PD-L1 expression on circulating tumor cells (CTCs) in patients with advanced urothelial carcinoma (UC). Oncoimmunology. Taylor & Francis; 2020;9:1738798.

41. Wang Y, Kim TH, Fouladdel S, Zhang Z, Soni P, Qin A, et al. PD-L1 Expression in Circulating Tumor Cells Increases during Radio(chemo)therapy and Indicates Poor Prognosis in Non-small Cell Lung Cancer. Sci Rep. 2019;9:566.

42. Mansour FA, Al-Mazrou A, Al-Mohanna F, Al-Alwan M, Ghebeh H. PD-L1 is overexpressed on breast cancer stem cells through notch3/mTOR axis. Oncoimmunology. 2020;9:1729299.

43. Yang J, Antin P, Berx G, Blanpain C, Brabletz T, Bronner M, et al. Guidelines and definitions for research on epithelial–mesenchymal transition. Nat Rev Mol Cell Biol. 2020;21:341–52.

44. Foroutan M, Bhuva DD, Lyu R, Horan K, Cursons J, Davis MJ. Single sample scoring of molecular phenotypes. BMC Bioinformatics. 2018;19:404.

45. Chakraborty P, George JT, Tripathi S, Levine H, Jolly MK. Comparative Study of Transcriptomics-Based Scoring Metrics for the Epithelial-Hybrid-Mesenchymal Spectrum. Front Bioeng Biotechnol. 2020;8:220.

46. Wang J, Zhang K, Xu L, Wang E. Quantifying the Waddington landscape and biological paths for development and differentiation. Proc Natl Acad Sci U S A. 2011;108:8257–62.

47. Subramanian A, Tamayo P, Mootha VK, Mukherjee S, Ebert BL, Gillette MA, et al. Gene set enrichment analysis: A knowledge-based approach for interpreting genome-wide expression profiles. Proc Natl Acad Sci U S A. 2005;102:15545–50.

48. Aibar S, González-Blas CB, Moerman T, Huynh-Thu VA, Imrichova H, Hulselmans G, et al. SCENIC: Single-cell regulatory network inference and clustering. Nat Methods. 2017;14:1083–6.

