## Supplementary Information for "Immunosuppressive traits of the hybrid epithelial/mesenchymal phenotype"

### Supplementary Text: Description of the ODE model

The chemical rate equation given below is a generic representation of the rate of change of the expression or level of each element (node).

$$\dot{E} = g_E * H^s(B,B_0,n,\lambda_{B,E}) - k_E * E \quad \text{(S1)}$$

Where $g_E$ is the basal production rate of $E$ and $k_E$ is the innate degradation rate of $E$, where $E$ represents expression level of a particular element (node). $H^S$ is the shifted Hill function, representing each interaction or regulatory term in the gene regulatory network between a pair of nodes, where $H^S$ represents interaction of node B affecting the production of node $E$ is defined as follows:

$$H^s(B,\lambda) = H^-(B) + \lambda H^+(B) \quad \text{(S2)}$$

$$H^-(B) = \frac{1}{(1 + \left(\frac{B}{B_0}\right)^{nB})}$$

$$H^+(B) = 1 - H^-(B)$$

$B_o$ = threshold value for that interaction,

n = cooperativity for that interaction,

λ = fold change from the basal synthesis rate of E due to B.

Hence, λ > 1 for activators and λ < 1 for inhibitors.

$$\frac{d\text{ZEB1}}{dt} = \text{g}_{\text{ZEB1}} * \text{H}^s(\text{miR200}, \lambda_{\text{miR200,ZEB1}}) * \text{H}^s(\text{ZEB1}, \lambda_{\text{ZEB1,ZEB1}}) * \text{H}^s(\text{CDH1}, \lambda_{\text{CDH1,ZEB1}}) * \text{H}^s(\text{SLUG}, \lambda_{\text{SLUG,ZEB1}}) - k_{\text{ZEB1}} * \text{ZEB1} \quad \text{(S3)}$$

$$\frac{d\text{miR200}}{dt} = \text{g}_{\text{miR200}} * \text{H}^s(\text{ZEB1}, \lambda_{\text{ZEB1,miR200}}) * \text{H}^s(\text{SLUG}, \lambda_{\text{SLUG,miR200}}) - k_{\text{miR200}} * \text{miR200} \quad \text{(S4)}$$

$$\frac{d\text{PDL1}}{dt} = \text{g}_{\text{PDL1}} * \text{H}^s(\text{PDL1}, \lambda_{\text{miR200,PDL1}}) - k_{\text{PDL1}} * \text{PDL1} \quad \text{(S5)}$$

$$\frac{d\text{CDH1}}{dt} = \text{g}_{\text{CDH1}} * \text{H}^s(\text{PDL1}, \lambda_{\text{PDL1,CDH1}}) * \text{H}^s(\text{ZEB1}, \lambda_{\text{ZEB1,CDH1}}) * \text{H}^s(\text{SLUG}, \lambda_{\text{SLUG,CDH1}}) - k_{\text{CDH1}} * \text{CDH1} \quad \text{(S6)}$$

$$\frac{d\text{SLUG}}{dt} = \text{g}_{\text{SLUG}} * \text{H}^s(\text{miR200}, \lambda_{\text{miR200,SLUG}}) * \text{H}^s(\text{SLUG}, \lambda_{\text{SLUG,SLUG}}) - k_{\text{SLUG}} * \text{SLUG} \quad \text{(S7)}$$

### Supplementary Tables:

| Wcoeff | ZeB1 | miR200 | SLUG | CDH1 | PDL1 |
|---|---|---|---|---|---|
| **PC1(84.44%)** | 0.46487 | -0.146355 | -0.34263 | -0.10234 | 0.79661 |
| **PC2(7.36%)** | -0.45943 | 0.21006 | 0.12549 | 0.72337 | 0.45361 |

***Table S1**: Contributions of the various node to the principal component axes PC-1 and PC-2 (**Fig S1A**).*

| Tissue | Number of cell lines | Spearman's correlation | log10(P-value) |
|---|---|---|---|
| **PROSTATE** | 8 | 0.714285714 | 1.332283447 |
| **LIVER** | 28 | 0.695675972 | 4.402583271 |
| **BILIARYTRACT** | 8 | 0.69047619 | 1.236644508 |
| **SOFTTISSUE** | 21 | 0.677922078 | 3.135173547 |
| **BREAST** | 59 | 0.654997078 | 7.739785255 |
| **PLEURA** | 11 | 0.636363636 | 1.452385494 |
| **KIDNEY** | 36 | 0.628828829 | 4.397764365 |
| **AUTONOMICGANGLIA** | 17 | 0.593137255 | 1.917750607 |
| **LUNG** | 187 | 0.515031619 | 13.33278092 |
| **LARGEINTESTINE** | 61 | 0.495769434 | 4.314209954 |
| **BONE** | 29 | 0.44137931 | 1.781603994 |
| **OVARY** | 52 | 0.433791514 | 2.881139725 |
| **ENDOMETRIUM** | 27 | 0.415140415 | 1.504534345 |
| **URINARYTRACT** | 27 | 0.368131868 | 1.230258399 |
| **HAEMATOPOIETICANDLYMPHOIDTISE** | 180 | 0.342856261 | 5.609133823 |
| **STOMACH** | 38 | 0.333406281 | 1.389322095 |
| **PANCREAS** | 44 | 0.286257928 | 1.224801918 |
| **CENTRALNERVOUSSYSTEM** | 69 | 0.242053343 | 1.34592534 |
| **SKIN** | 62 | 0.143763693 | 0.57683202 |
| **OESOPHAGUS** | 26 | 0.140512821 | 0.306655414 |
| **THYROID** | 12 | 0.048951049 | 0.055557512 |
| **UPPERAERODIGESTIVETRACT** | 32 | 0.02016129 | 0.039630176 |

***Table S2**: Tissue specific CCLE spearman's correlation and p values of PD-L1 expression levels with ssGSEA scores of Hallmark EMT gene set (**Fig 1E** – right panel).*

| S. No | Interactions | Reference |
|---|---|---|
| **1** | ZEB1, miR200 mutual inhibition | (1) |
| **2** | ZEB1 self-activation | (2) |
| **3** | miR200 downregulate PDL1 | (3) |
| **4** | PD-L1 downregulate CDH1 | (4) |
| **5** | ZEB1, CDH1 mutual inhibition | (5–7) |
| **6** | SLUG, miR200 mutual inhibition | (8) |
| **7** | SLUG upregulate ZEB1 | (9) |
| **8** | SLUG downregulate CDH1 | (10) |
| **9** | SLUG self-activation | (11) |
| **10** | OCT4 self-activation | (12) |
| **11** | OCT4 upregulates miR-200 | (13) |
| **12** | OCT4 upregulates SLUG | (14) |
| **13** | PD-L1 upregulate OCT4 | (15) |
| **14** | LIN28 upregulate OCT4 | (16) |
| **15** | OCT4, miR-145 mutual inhibition | (13) |
| **16** | let7 self-activation | (16) |
| **17** | let7 downregulate PD-L1 | (17) |
| **18** | let7 downregulate ZEB1 | (16) |
| **19** | let7, LIN28 mutual inhibition | (16) |
| **20** | LIN28 self-activation | (16) |
| **21** | miR-200 downregulates LIN28 | (16) |
| **22** | miR-145, ZEB1 mutual inhibition | (18) |
| **23** | miR-145, SLUG mutual inhibition | (19, 20) |
| **24** | SLUG, ERα66 mutual inhibition | (21) |
| **25** | ZEB1 downregulate ERα66 | (21) |
| **26** | ERα66 self-activation | (21) |
| **27** | ERα66 downregulate ERα36 | (21) |
| **28** | ERα66 downregulate PD-L1 | (22) |
| **29** | ERα36 upregulate ZEB1 | (21) |

***Table S3**: Interactions for core GRN (**Fig 1A**; S. No 1-11), Interactions for stemness circuit (**Fig S4C**; S. No 1-23), Interactions for drug resistance circuit (**Fig 4A**; S. No 1-11 & S. No 24-29).*

| Parameters | Minimum - Maximum (Uniform Values) |
|---|---|
| **Maximum production rate (*g*)** | 1-100 |
| **Degradation rate (*k*)** | 0.1-1 |
| **Fold change (*λ*)** | 1-100 |
| **Threshold ($B_o$)** | The ranges depend on the inward regulations, which are estimated by a Monte Carlo simulation. |
| **Hill coefficient (*n*)** | 1-6 |

***Table S4**: Ranges of the parameters within which RACIPE randomly samples.*

| Parameter | Value | Parameter | Value |
|---|---|---|---|
| Prod_of_ZEB1 | 51.20536 | Inh_of_SLUGTomiR200 | 0.01051 |
| Prod_of_miR200 | 73.74496 | Trd_of_miR200ToPDL1 | 9.788561 |
| Prod_of_PDL1 | 43.96804 | Num_of_miR200ToPDL1 | 6 |
| Prod_of_CDH1 | 89.20117 | Inh_of_miR200ToPDL1 | 0.01356 |
| Prod_of_SLUG | 29.23346 | Trd_of_PDL1ToCDH1 | 34.06882 |
| Deg_of_ZEB1 | 0.62873 | Num_of_PDL1ToCDH1 | 2 |
| Deg_of_miR200 | 0.861129 | Inh_of_PDL1ToCDH1 | 0.010248 |
| Deg_of_PDL1 | 0.837638 | Trd_of_ZEB1ToCDH1 | 0.80368 |
| Deg_of_CDH1 | 0.192489 | Num_of_ZEB1ToCDH1 | 3 |
| Deg_of_SLUG | 0.549676 | Inh_of_ZEB1ToCDH1 | 0.041006 |
| Trd_of_miR200ToZEB1 | 6.155112 | Trd_of_SLUGToCDH1 | 11.37985 |
| Num_of_miR200ToZEB1 | 2 | Num_of_SLUGToCDH1 | 6 |
| Inh_of_miR200ToZEB1 | 0.014161 | Inh_of_SLUGToCDH1 | 0.06103 |
| Trd_of_ZEB1ToZEB1 | 0.662827 | Trd_of_miR200ToSLUG | 8.200188 |
| Num_of_ZEB1ToZEB1 | 2 | Num_of_miR200ToSLUG | 3 |
| Act_of_ZEB1ToZEB1 | 54.05437 | Inh_of_miR200ToSLUG | 0.16813 |
| Trd_of_CDH1ToZEB1 | 0.547427 | Trd_of_SLUGToSLUG | 13.8752 |
| Num_of_CDH1ToZEB1 | 2 | Num_of_SLUGToSLUG | 3 |
| Inh_of_CDH1ToZEB1 | 0.01252 | Act_of_SLUGToSLUG | 25.64712 |
| Trd_of_SLUGToZEB1 | 10.8639 | Num_of_ZEB1TomiR200 | 1 |
| Num_of_SLUGToZEB1 | 5 | Inh_of_ZEB1TomiR200 | 0.018446 |
| Act_of_SLUGToZEB1 | 72.00952 | Trd_of_SLUGTomiR200 | 5.575322 |
| Trd_of_ZEB1TomiR200 | 1.573366 | Num_of_SLUGTomiR200 | 6 |

***Table S5**: Parameter values for generation of probability landscape and steady state plot (**Fig 2A, B**). Here, production terms are represented in green color, degradation terms are represented in orange color, Hill coefficients represented in cyan color, Threshold terms represented in pink color and fold change represented in white color.*

| Parameter | Value | Parameter | Value |
|---|---|---|---|
| G_ZEB1 | 62.16564 | FC_ZEB1_ZEB1 | 68.19913 |
| G_miR200 | 74.63792 | FC_miR200_ZEB1 | 91.63278 |
| G_PDL1 | 44.80946 | FC_CDH1_ZEB1 | 76.93204 |
| G_CDH1 | 80.99668 | FC_SLUG_ZEB1 | 97.67022 |
| G_SLUG | 32.3487 | FC_ZEB1_miR200 | 67.84465 |
| K_ZEB1 | 0.529706 | FC_SLUG_miR200 | 67.49304 |
| K_miR200 | 0.330501 | FC_miR200_PDL1 | 68.41486 |
| K_PDL1 | 0.928506 | FC_ZEB1_CDH1 | 94.1392 |
| K_CDH1 | 0.88837 | FC_PDL1_CDH1 | 38.9354 |
| K_SLUG | 0.518909 | FC_SLUG_CDH1 | 73.84352 |
| TH_ZEB1_ZEB1 | 0.902465 | FC_miR200_SLUG | 56.15421 |
| TH_miR200_ZEB1 | 4.090531 | FC_SLUG_SLUG | 72.962 |
| TH_CDH1_ZEB1 | 0.623593 | N_miR200_ZEB1 | 4 |
| TH_SLUG_ZEB1 | 2.210129 | N_CDH1_ZEB1 | 3 |
| TH_ZEB1_miR200 | 0.814146 | N_SLUG_ZEB1 | 2 |
| TH_SLUG_miR200 | 6.22973 | N_ZEB1_miR200 | 5 |
| TH_miR200_PDL1 | 3.811111 | N_SLUG_miR200 | 4 |
| TH_ZEB1_CDH1 | 0.762967 | N_miR200_PDL1 | 2 |
| TH_PDL1_CDH1 | 9.033578 | N_ZEB1_CDH1 | 1 |
| TH_SLUG_CDH1 | 12.66515 | N_PDL1_CDH1 | 3 |
| TH_miR200_SLUG | 8.263242 | N_SLUG_CDH1 | 1 |
| TH_SLUG_SLUG | 8.61641 | N_miR200_SLUG | 5 |
| N_ZEB1_ZEB1 | 2 | N_SLUG_SLUG | 1 |

***Table S6****: Basic parameter values. The parameters were adopted from sRACIPE (****Fig. 2C****), which generate random set of parameters and to simulate the system with a fixed amount of noise. In the table production terms are represented in green color, degradation terms are represented in orange color, Hill coefficients represented in cyan color, Threshold terms represented in pink color and fold change represented in white color.*

### References


1. U. Burk, *et al.*, A reciprocal repression between ZEB1 and members of the miR-200 family promotes EMT and invasion in cancer cells. *EMBO reports* **9**, 582–589 (2008).

2. L. Hill, G. Browne, E. Tulchinsky, ZEB/miR-200 feedback loop: At the crossroads of signal transduction in cancer. *International Journal of Cancer* **132**, 745–754 (2013).

3. L. Chen, *et al.*, Metastasis is regulated via microRNA-200/ZEB1 axis control of tumour cell PD-L1 expression and intratumoral immunosuppression. *Nature Communications* **5**, 5241 (2014).

4. W. Yu, *et al.*, PD-L1 promotes tumor growth and progression by activating WIP and β-catenin signaling pathways and predicts poor prognosis in lung cancer. *Cell Death & Disease* **11**, 506 (2020).

5. A. B. Singh, *et al.*, Claudin-1 Up-regulates the Repressor ZEB-1 to Inhibit E-Cadherin Expression in Colon Cancer Cells. *Gastroenterology* **141**, 2140–2153 (2011).

6. Y.-F. Tan, *et al.*, β-catenin-coordinated lncRNA MALAT1 up-regulation of ZEB-1 could enhance the telomerase activity in HGF-mediated differentiation of bone marrow mesenchymal stem cells into hepatocytes. *Pathology - Research and Practice* **215**, 546–554 (2019).

7. S. Orsulic, O. Huber, H. Aberle, S. Arnold, R. Kemler, E-cadherin binding prevents beta-catenin nuclear localization and beta-catenin/LEF-1-mediated transactivation. *Journal of Cell Science* **112**, 1237–1245 (1999).

8. Y.-N. Liu, *et al.*, MiR-1 and miR-200 inhibit EMT via Slug-dependent and tumorigenesis via Slug-independent mechanisms. *Oncogene* **32**, 296–306 (2013).

9. C. Wels, S. Joshi, P. Koefinger, H. Bergler, H. Schaider, Transcriptional activation of ZEB1 by Slug leads to cooperative regulation of the epithelial-mesenchymal transition-like phenotype in melanoma. *The Journal of investigative dermatology* **131**, 1877–1885 (2011).

10. K. M. Hajra, D. Y.-S. Chen, E. R. Fearon, The SLUG zinc-finger protein represses E-cadherin in breast cancer. *Cancer research* **62**, 1613–1618 (2002).

11. B. Kumar, *et al.*, Auto-regulation of Slug mediates its activity during epithelial to mesenchymal transition. *Biochimica et biophysica acta* **1849**, 1209–1218 (2015).

12. G. Shi, Y. Jin, Role of Oct4 in maintaining and regaining stem cell pluripotency. *Stem Cell Research & Therapy* **1**, 39 (2010).

13. M. K. Jolly, *et al.*, Stability of the hybrid epithelial/mesenchymal phenotype. *Oncotarget; Vol 7, No 19* (2016).

14. G. Mandal, *et al.*, Heterodimer formation by Oct4 and Smad3 differentially regulates epithelial-to-mesenchymal transition-associated factors in breast cancer progression. *Biochimica et Biophysica Acta (BBA) - Molecular Basis of Disease* **1864**, 2053–2066 (2018).

15. S. Almozyan, *et al.*, PD-L1 promotes OCT4 and Nanog expression in breast cancer stem cells by sustaining PI3K/AKT pathway activation. *International journal of cancer* **141**, 1402–1412 (2017).

16. M. K. Jolly, *et al.*, Coupling the modules of EMT and stemness: A tunable "stemness window" model. *Oncotarget* **6**, 25161–25174 (2015).

17. Y. Chen, *et al.*, LIN28/<em>let-7</em>/PD-L1 Pathway as a Target for Cancer Immunotherapy. *Cancer Immunology Research* **7**, 487 LP – 497 (2019).

18. K. Hari, *et al.*, Identifying inhibitors of epithelial–mesenchymal plasticity using a network topology-based approach. *npj Systems Biology and Applications* **6**, 15 (2020).

19. L.-L. Mei, *et al.*, miR-145-5p Suppresses Tumor Cell Migration, Invasion and Epithelial to Mesenchymal Transition by Regulating the Sp1/NF-κB Signaling Pathway in Esophageal Squamous Cell Carcinoma. *International journal of molecular sciences* **18**, 1833 (2017).

20. V. J. Findlay, *et al.*, SNAI2 modulates colorectal cancer 5-fluorouracil sensitivity through miR145 repression. *Molecular cancer therapeutics* **13**, 2713–2726 (2014).

21. S. Sahoo, *et al.*, A mechanistic model captures the emergence and implications of non-genetic heterogeneity and reversible drug resistance in ER+ breast cancer cells. *NAR Cancer* **3** (2021).

22. L. Liu, *et al.*, ERα is a negative regulator of PD-L1 gene transcription in breast cancer. *Biochemical and biophysical research communications* **505**, 157–161 (2018).
